# Individual Differences in Policy Precision: Links to Suicidal Ideation and Network Dynamics

**DOI:** 10.1101/2024.10.15.618415

**Authors:** Dayoung Yoon, Jaejoong Kim, Do Hyun Kim, Dong Woo Shin, Su Hyun Bong, Jaewon Kim, Hae-Jeong Park, Hong Jin Jeon, Bumseok Jeong

## Abstract

Behavioural modelling of decision-making processes has advanced our understanding of impairments associated with various psychiatric conditions. However, as research increasingly prioritises the development of models that best explain observed behaviour, the question of whether these behaviours stem from biologically plausible brain functions has often been overlooked. To address this gap, we developed a probabilistic two-armed bandit task model based on the active inference framework and compared its performance to established reinforcement learning (RL) models. Our model demonstrated superior explanatory power in capturing individual variability in choice behaviour. A key parameter in our model, policy precision—analogous to the temperature parameter in RL models—is also optimised based on previous outcomes. This optimisation accounts for the balance between model-free (MF) and model-based (MB) decision-making strategies. Notably, incorporating the rate of change in policy precision enhanced the model’s ability to explain brain network dynamics and their inter-subject correlations. Specifically, we observed a positive correlation with default mode network dominance and a negative correlation with dorsal attention and frontoparietal network-dominant states. These opposing network patterns suggest a cooperative relationship, as evidenced by correlations between state transitions and behavioural parameters. This transition may represent a neural mechanism underlying MB-MF arbitration, which appears to be disrupted by prolonged activation of another state characterised by heightened ventral attention network activity and increased inter-network connectivity. Finally, we found that reduced prior policy precision in loss-related context is associated with suicidal ideation in individuals with major depressive disorders.

**Highlights:**

- The AIF model explains pronounced individual behavioural variability.
- Neural signals are better explained by changes in policy precision.
- The anti-correlation can be explained from the perspective of the MB-MF arbitration.
- The AIF model better explains the HAM-D score.
- The AIF model can discriminate suicidal ideation in MDD with a loss task.

## Introduction

Computational psychiatry has taken on the challenge of bridging the gap between mental phenomena and their assumed origins in brain function [1, 2]. Central to this effort has been the development of mathematical models. Owing to the intricate cognitive processes involved in decision-making, such as evaluating evidence for potential outcomes and option selection, decision-making has become a means to explain psychiatric symptoms [3, 4]. Reinforcement learning (RL) has facilitated decision-making and the development of various models augmented with additional parameters to more accurately explain the agent’s decision [5–8]. Building on previous studies linking major depressive disorder (MDD) to abnormalities in the brain’s reward circuitry, RL has been widely used to explain decision-making deficits in patients with MDD [3, 9, 10]. There are various decision-making tasks, one of the simplest forms of which is the probabilistic two-armed bandit task. Owing to its simplicity, it has been neglected since it is considered insufficient for studying complex cognitive processes, such as exploration [11, 12]. However, even in this simple task, individuals display diverse behavioural patterns likely rooted in variations in neural states. Although behavioural models are intended to link these neural and behavioural patterns, most account for this behavioural diversity merely by randomness in action probabilities and exclude neural state diversity from consideration. This likely stems from the fact that most behavioural models have been developed to explain relatively low-dimensional behavioural patterns, without explaining the high-dimensional diversity of neural states. To address this limitation, we developed a new model inspired by the principles of the active inference framework (AIF) and determined whether the behaviourally relevant signals central to this model can account for variations in brain imaging data using a general linear model (GLM) in two ways. AIF is a unified theory of action and perception that attempts to elucidate decision-making processes by minimising a singular objective function, free energy [13]. Free energy measures the discrepancy between an agent’s internal model (its beliefs about the world) and the actual sensory data it receives, and, in AIF, serves as a proxy for surprise (unexpectedness), which cannot be directly computed by agents.

The RL model assumes that people develop expected values (EV) for each choice option that represent the reward (or punishment) they expect to receive following each choice. In this process of learning, a Rescorla-Wagner model is generally implemented, which uses a fixed learning rate parameter for updating expected values [14].

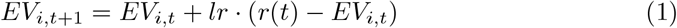

Choices are made according to the probability calculated from EVs, which are updated and learned through observations, using the softmax action selection rule. The degree to which the option with the highest EV is chosen is determined by *γ* referred to as the ‘decision temperature’.

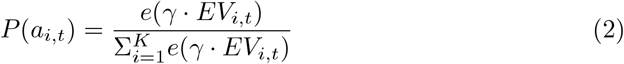

However, this method determines the probability of selecting each option solely based on its EVs, which cannot capture the trial-by-trial variability in individual behaviour. In other words, it does not adequately account for within-subject fluctuations that unfold as the task progresses. The decision temperature parameter can partially compensate for this limitation by modulating the influence of EVs; when choice inconsistency increases, the estimated temperature decreases. This results in individuals with more inconsistent behaviour being interpreted as placing less weight on value-based decisions, a feature often linked to impulsivity in decision-making [15]. In this sense, this model is well-suited to explain overall observed behavioural data. However, the model falls short when answering questions like, “Why did the person make that specific choice at that specific moment?”. This limitation is especially apparent when explaining moments of increased choice variability across individuals. As shown in 3A, B, the entropy of choices across participants is highest at the beginning of the task and decreases after approximately 10 trials, suggesting that most participants learned which bandit is better by that point. Interestingly, there are certain trials later in the task where entropy increases again. The initial rise in entropy can be explained by the absence of knowledge, where EVs are close to 0.5, leading to action probabilities that approximate the chance level. However, the later increases in entropy tend to follow surprising events, such as consecutive negative outcomes from selecting the better bandit.

The basic model in Equations (1-2) attributes such behaviour to an overall increase in randomness as captured by decreased EV of better bandit consecutive negative outcomes. However, the entropy in these trials is not high enough to suggest total uncertainty, since it is in the early phase of the task. This implies the model lacks an important explanatory term beyond EVs to account for action selection in these moments. Such a term must capture how individuals respond to outcomes from the previous trial. A well-known strategy that directly reflects this is the win-stay-lose-shift (WSLS) strategy, which is entirely outcome-driven [16–18]. Prior research has suggested that both methods are can be utilised in decision-making; some studies have attempted to integrate both approaches [7, 8]. The WSLS-RL model, for example, combines both probabilities—those derived from EV and WSLS strategy—by weighting them with a parameter *w*.

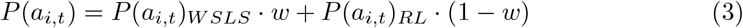

In this model, the parameter *w* and the probability of lose-shift (*P*_*WSLS*_) correspond to the parameters that can explain the choice variability in specific trials. However, this method relies heavily on additional parameters and cannot explain the mechanism involved in the exchange between both strategies according to the state context.

Meanwhile, two aspects of AIF allow for a flexible model to address this limitation. First, rather than using EV directly, the expected free energy (EFE) is applied as the input to calculate choice probability. EFE represents the distance between the probability distribution of this EV and each agent’s preference distribution for EV (Equation 7). This approach accounts for utility differences [5] and captures variations in EV precision, which can influence choice. For instance, when the mean of EV for two options is identical, calculating EFE with higher precision in one case and lower in another yields a smaller EFE difference between the two options in the latter case, making the choice probability more similar to chance (Figure1A). Second, the policy precision parameter (*γ* in Equation 9), which corresponds to the decision temperature in a softmax function, modulates the effect of EFE on action probability. This parameter is updated as the differences between EFE and free energy (Equation 13, Figure 1B). Here, EFE reflects a model-based (MB) strategy in RL calculated based on accumulated evidence and preferences over future outcomes. In contrast, free energy corresponds to prediction error and represents a model-free (MF) strategy (the Discussion section provides further explanation regarding the MB–MF distinction).

**Fig. 1.**
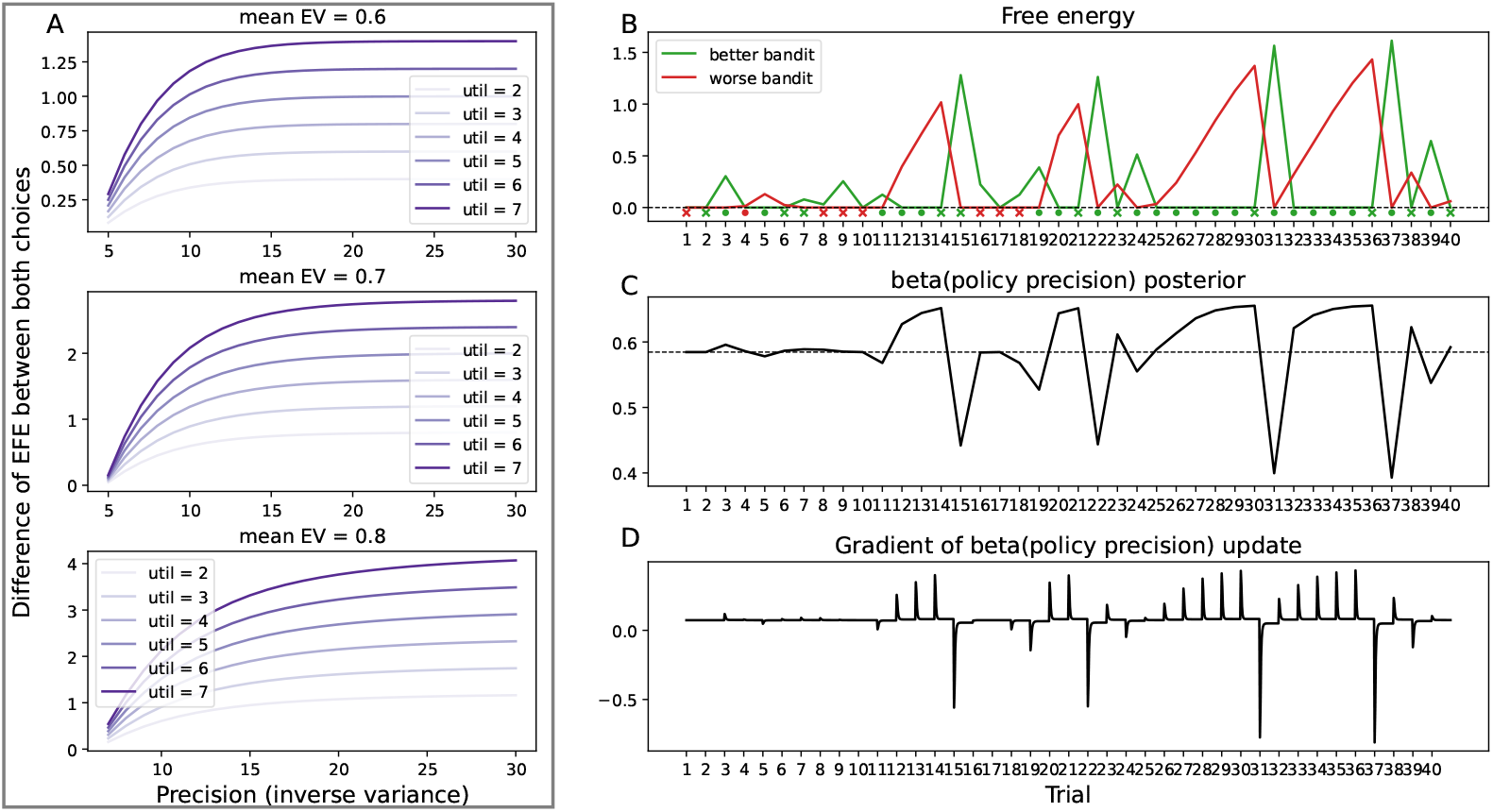
Simulated expected free energy and example estimated results of AIF model. (A) Differences in expected free energy values between both actions increase as the precision of the state increases, regardless of the expected value of the state. (B–D) The estimated results for one participant. (B) Free energy values for both actions in each trial. These values are generally higher during the later phase of the trials because the variance of the state distribution is lower. In this phase, if the outcomes differ from the prediction, the distribution’s distance to the predicted state is large. Conversely, free energy values before approximately the tenth trial are lower because the prediction precision is low; therefore, the prediction error is also low. Each bandit’s free energy tends to increase after it is selected and a losing outcome is observed. (C) Since the policy precision is also updated according to the free energy, updated posterior values tend to follow the worse bandit’s free energy while opposing the free energy of the better bandit. (D) While policy precision is updated, the rate of change is hypothesised to represent neural discharge.

The update of policy precision allows the probability of action selection to be more influenced by either EFE or free energy. Thus, without additional weighting parameters, the model can modulate the alternation between both strategies as a function of state uncertainty. This aligns with previous findings indicating that uncertainty influences the arbitration process between MF and MB, such that high levels of uncertainty lead participants to rely more on the MF strategy [19]. A detailed explanation of the model is provided in the Materials and Methods section 4.

We propose a new behavioural model based on AIF for the probabilistic two-armed bandit task and demonstrate its advantages in explaining inter-subject variability in choice behaviour. We argue that these advantages stem from the model’s ability to capture the notion that confidence in decision making, as reflected in decision variability, is influenced by disconfirmatory evidence, over and above the change in pragmatic action value that such evidence induces. By incorporating this additional influence, the model can better explain within-subject variability that emerges throughout the task. To evaluate how well the parameters estimated by each model capture inter-subject variability, we computed the cross-entropy of individual choices across trials. We calculated the mutual information between cross-entropy and the behavioural parameters, with higher values indicating better explanatory power [20]. In addition, the validity of the policy precision signal (rate of change in policy precision, Figure 1D) is confirmed through two experiments: one involving the analysis of neural signal data and another examining whether behaviour parameters provide better explanatory power in describing traits of depressive patients.

In Experiment 1, using neural signal data obtained via task-functional magnetic resonance imaging (fMRI), we used two methods to confirm whether the policy precision signal is related to neural activity (Figure 4). First, we conducted a conventional GLM approach, where models are generated by including the stimuli of behavioural tasks as regressors. The second method addresses the limitations of the conventional GLM by using inter-subject correlation [21]. We determined whether the variance in brain imaging data across individuals could be explained by the variance in behavioural signals and, similarly, whether adding policy precision signal improves model performance measured by Akaike Information Criterion (AIC).

In Experiment 2, we conducted the decision-making task involving a population that included patients diagnosed with MDD and healthy controls (HC). In patients with MDD, the risk of suicide is nearly 20 times higher than that in the general population [22]. According to the Diagnostic and Statistical Manual of Mental Disorders, 5th edition (DSM-5), “impairment in decision-making” is an important criterion for diagnosing MDD as “indecisiveness”. Previous studies have analysed patients with MDD with serious suicidal behaviour and found that their ability to learn and choose actions leading to rewarding outcomes is impaired [23–25]. Although most studies on suicide have shown impaired decision-making and its associated neural correlates within the reward domain, impairments in the loss domain have rarely been investigated, especially in young patients with MDD and suicidal risk. Furthermore, a review by [3] highlighted inconsistent findings across studies employing reinforcement learning tasks in the reward and punishment domains. In this study, we aim to determine whether estimated parameters from a decision-making task involving reward and loss domains can distinguish MDD subgroups categorised by suicidal ideation, and whether the newly proposed AIF model offers improved adaptability for this purpose.

## 2 Results

### 2.1 Model construction and the results of parameter recovery and model comparison

We constructed 16 models, comprising 4 learning models combined with 4 choice models. One of the learning models utilised scalar state value explained in Equation 1 (Rescorla-Wagner in Supplementary Table. 1), whereas the others utilised either beta or normal distributions for state value. The latter corresponds to the Kalman-filter model [26]. With beta distribution, the ‘decay’ parameter *ρ* is added during belief updating (Equation 4) that prevents beliefs from becoming quickly inflexible. This term can be a fixed value across the trials or changed based on the Bayes-factor surprise, a ratio between the subjective probability of an observation under the current beliefs and prior beliefs [27]. The four choice models included two previous models of the softmax rule (*sft* in Supplementary Table. 1) and the WSLS-RL (*wsls* in Supplementary Table. 1) model. The other two models used EFE, referred to as the *efeX* and *efegc* models. The *efeX* model excluded the process of policy precision update, making it similar to the softmax model, whereas the *efegc* model resembled the WSLS-RL model. The parameter compositions and recovery results of each model are described in Supplementary Table. 1 with descriptions of each parameter. The model construction is detailed in the Materials and methods section 4. Parameter estimation was accomplished using the variational Bayes method [28], and the approximations of log model evidence were subjected to random-effect Bayesian model selection [29]. First, we grouped 16 models into 4 ‘families’ by learning models [30]. This method yields family-specific exceedance probabilities, representing the probability that the learning model has a higher frequency than the other included models on the group level. The beta distribution decay modulated by the surprise model outperformed the others in loss and reward tasks (Figure 2A, B). Parameter recovery was examined with simulation data generated from the distributions of each estimated parameter. Except for both beta distribution models, other models showed poor parameter recovery performance in at least one action model combination (Supplementary Table. 1). Based on these results, we selected the models incorporating beta distribution decay modulated by surprise for subsequent analyses. Then we compared the four action models in that learning model family. The model with the softmax rule was best for loss and reward tasks (Figure 2C, D).

**Fig. 2.**
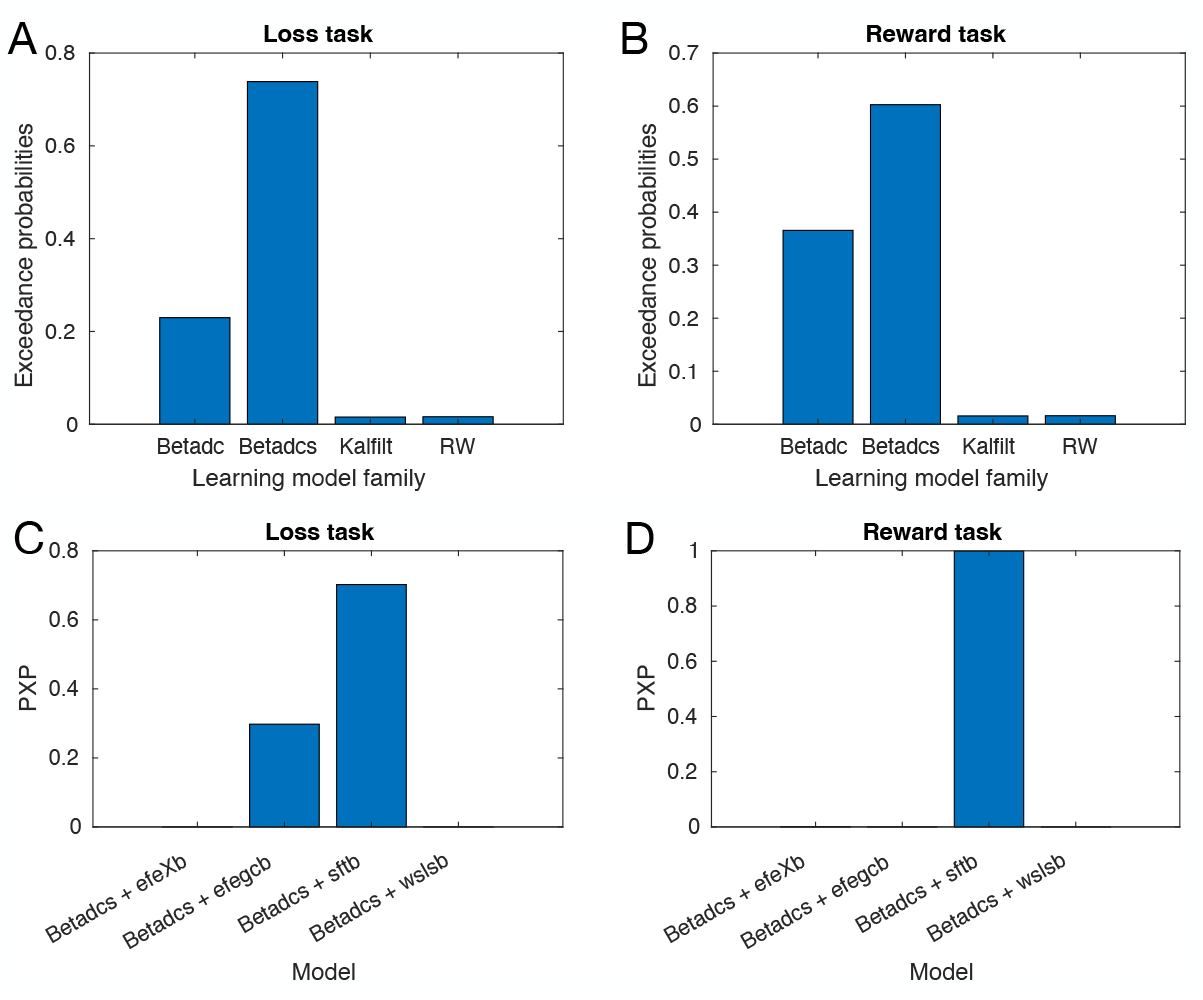
Model comparison results. (A, B) Result of random-effect Bayesian model selection performed with grouping 16 models with 4 learning models in both tasks. (C,D) Result of random-effect Bayesian model selection among the action models within the beta distribution with updated decay learning model in both tasks. PXP; protected exceedance probability, Betadc; beta distribution with fixed decay model, Betadcs; beta distribution with updated decay model, Kalfilt; Kalman filter model, RW; Rescorla-Wagner model, efeX: active inference model without the process of policy precision update, efegc; active inference model including the process of policy precision update, sft; softmax model, wsls; win-stay-lose-shift model

### 2.2 Choice variability across individuals explained by deviations of behavioural parameter

Model comparison results showed that the softmax rule was best for loss and reward tasks (Figure 2C, D). However, this method had critical limitations in the two-armed bandit task: one choice was better, and the task’s objective was to learn which was better. As a result, the better bandit was selected far more frequently than the other. In such cases, models that estimate action probabilities overly favouring the better bandit are more likely to be evaluated as optimal, even if they lack a balanced perspective. For instance, when the action probability for the better bandit (p) is estimated as a large value close to 1, the selection probability for the other bandit decreases to (1 - p). Since the worse bandit accounts for only a small proportion of the total trials (as it is rarely selected), the sum of the log probabilities for the selected bandit can still yield a large final log-likelihood value. This creates a problem as it generates a bias in model evaluation favouring models that heavily prioritise the better bandit, even if they fail to represent the data comprehensively. In the case of the softmax model, simply increasing the decision temperature parameter, *β*_0_, results in action probabilities approaching 1 for the better bandit, thereby increasing the likelihood. As a consequence, the estimated parameter tends to converge to a high value that does not capture individual differences. In this way, the fitted parameters are limited in their ability to reflect between-subject variability in behaviour, thus failing to account for cognitive differences across individuals. In other words, the model’s strong performance in terms of likelihood does not guarantee that it optimally explains the underlying cognitive processes, especially in the infrequent but critical cases where choices deviate from the expected pattern in ways that vary across individuals.

Figure 3A, B illustrates the entropy of binary choices across the individuals. On the first trial, the selection variability was highest because there was no difference in value between bandits, suggesting that this early variance is driven by random selection. In contrast, the increase in variance observed following the tenth trial is attributed to a distinct mechanism related to the direct influence of the previous trial’s result, which is encoded as the lose-shift probability (*ls*) in the WSLS-RL model. The softmax model accounts for this by the increased selection randomness due to a temporary reduction in the value difference between two options, similar to what occurred during the early phase. The cross-entropy (CE) for each participant quantifies the extent to which their choice trajectory deviates from the group norm. It is computed as:

**Fig. 3.**
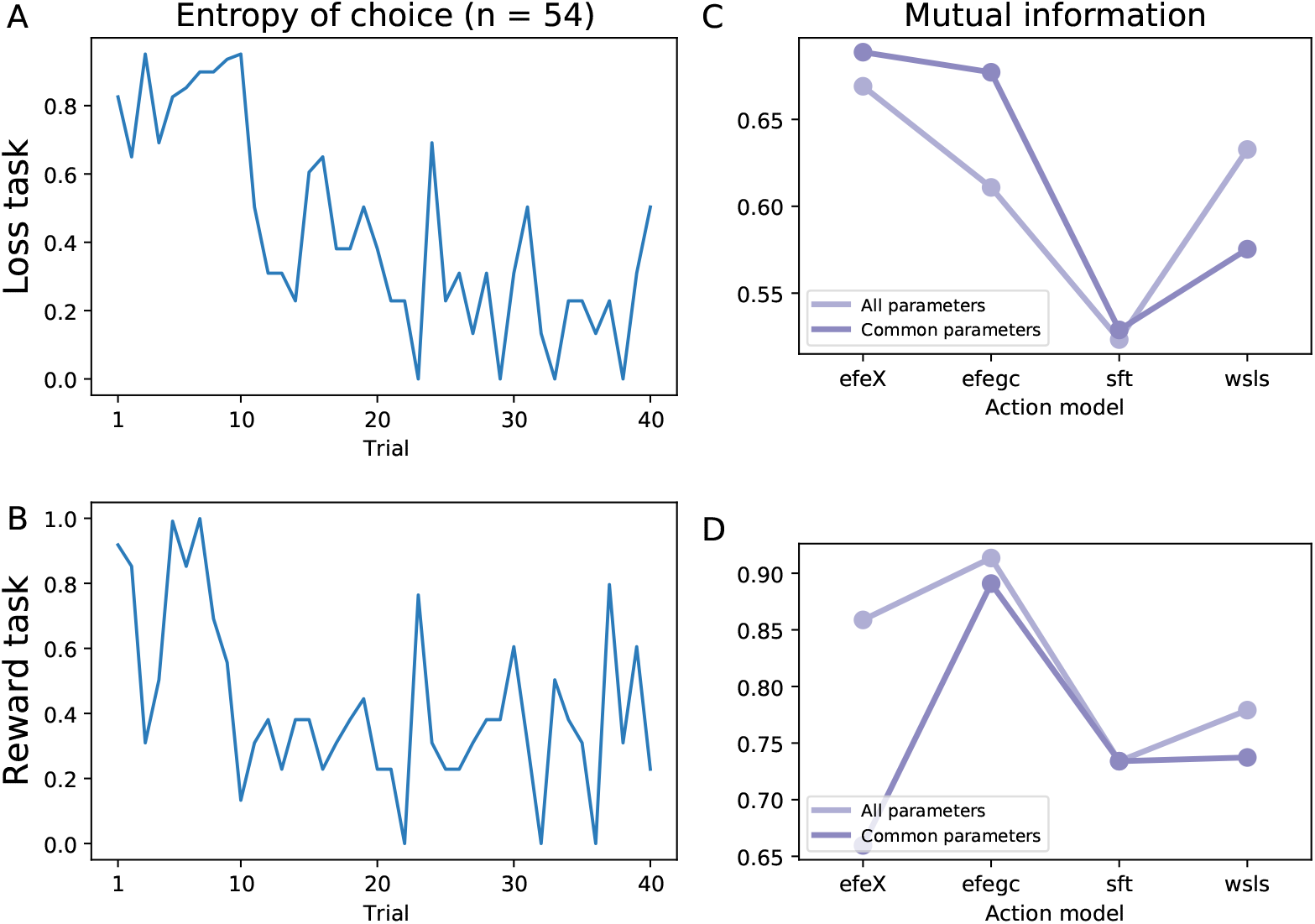
Entropy of choices across the participants and mutual information. (A, B)The entropy of choices in each trial out of 54 participants. (C, D) Mutual information between the cross-entropy of behaviour and the distance from the centre in the PCA space of behavioural parameters. Each of the four action models includes a different number of behavioural parameters—3, 3, 2, and 4, respectively. All models share the same learning model parameters, including *ρ*, and include *β*_0_ as a common parameter across action models.

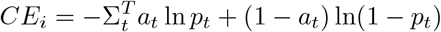

where *a*_*t*_ ∈ 0, 1 represents the choice made by the participant *i* on trial *t*, and *p*_*t*_ denotes the group-level probability of choosing option 1 on that trial. Close to 0.5, trial entropy is high, meaning that participants’ choices are relatively unpredictable and equally split. In such cases, individual differences contribute little to the CE, since either choice is plausible. Similarly, when *p*_*t*_ is near 0 or 1 (i.e., strong group consensus), the contribution to CE is minimal, as there is little behavioural variability. Importantly, the trials that contribute most to inter-individual differences in CE are those where *p*_*t*_ is moderately skewed—that is, not close to 0.5, yet not extreme. On such trials, a participant making a choice that deviates from the group (e.g., choosing 0 when *p*_*t*_ is high) incurs a larger increase in CE, reflecting more idiosyncratic decision-making. If these idiosyncratic decision-making patterns are reflected in behavioural parameters, then the CE values should be well predicted by those parameters. To test this, we performed principal component analysis (PCA) on the vector of behavioural parameters and computed each participant’s distance from the origin (i.e., the centre of the PCA space). We then examined the relationship between this distance and the participant’s CE value. To capture potential nonlinear associations between the two variables, we also computed their mutual information. Consistent with our expectations, individual differences in parameters estimated from the soft-max model accounted for less inter-subject behavioural variability than those from the other models (Figure 3C, D).

### 2.3 Neuroimaging analysis

Although model comparison based on model evidence indicated that the softmax model provided the best fit, behavioural parameters estimated from other models better captured inter-subject variability in behaviour. This improvement stems from the additional parameters, *ls* and *w*, in the WSLS-RL model and the inclusion of an additional process in *efegc* that optimises policy precision by modulating the dependency between MF and MB strategies. Notably, *efegc* does not introduce any additional parameters but includes an additional signal represented by the rate of change in policy precision. This point is further supported by the analysis conducted using only the shared parameters across models, *ρ* and *β*_0_, where the mutual information for the WSLS-RL model dropped substantially, comparable to the softmax model, suggesting that the improvement in WSLS-RL largely arises from its additional parameters (Figure 3C, D). To determine whether this policy precision signal is suitable for explaining brain function, we compared behavioural signals and neural signals across three levels: node, edge, and network. This approach captures regional brain activity and changes in connectivity between regions and alterations in large-scale networks critical to brain function (Figure 4). We estimated four key states using the graph Laplacian mixture model (GLMM) as described by [31]. A detailed description of the neural signal extraction process is provided in the Materials and Methods section4. Using the mean activation patterns, we grouped the regions into a priori resting-state networks (RSNs) based on [32] (Supplementary Figure. 1A-D). State 1 was positively correlated with activation of the visual (VIS) and somatomotor (SOM) networks, dorsal attention network (DAN), and ventral attention network (VAN) and negatively correlated with the default mode network (DMN) and frontoparietal control network (FPN) and limbic network (LIM). State 2 was positively correlated with the DMN and negatively correlated with State 1’s network. State 3 was positively correlated with the DAN and FPN and negatively correlated with the DMN and SOM network. State 4 shows generally low activations of most networks while those of the LIM and VAN are relatively high. We found that adding policy precision signals to the GLM yielded lower AIC values, which corresponds to better performance (Figure 4 I-N). Using beta coefficients from the best model, we performed one-sample t-test to identify which regional activations or interactions are related to the policy precision signals (Supplementary Table. 3). Across both tasks, similar regions included in DMN were positively correlated with policy precision signal (Figure 5), whereas regions included in the FPN and DAN showed negative correlations. These results are consistent with the results of the state level that occurrences of State 2 showed a positive correlation with policy precision signals (*p* = 0.009 for loss task, *p* = 0.021 for reward task) whereas State 3 exhibited a negative correlation (*p* = 9.77 × 10^*−*5^ for loss task, *p* = 0.013 for reward task). Using the occurrences of each state, we computed three of the state dynamics metrics for each participant: fractional occupancy, dwell time, and transition probability [33]. We then examined whether these metrics were associated with behavioural indicators using Pearson correlation analyses. The behavioural indicators were latent behavioural parameters estimated using the *betadcs* with *efegc* model, which yielded three parameters: prior of policy precision (*beta0*), volatility (*v*), and *util* (reward sensitivity or loss aversiveness). To derive a single summary measure, we applied PCA to the parameters. Due to the high correlation between *beta0* and *util* (r = 0.998), *util* was excluded from PCA to maintain balance between variables. Additionally, because the *v* reflects an opposite characteristic—higher values indicate reduced choice consistency and increased sensitivity to recent outcomes-and it was negatively correlated with the *beta0* (*r* = 0.72 and 0.79 for the loss and reward tasks, respectively), we applied PCA to the negative of the *v*. The first principal component explained 66.7%. This component was a weighted combination of the three parameters, with loadings of 0.52, 0.45, −0.51, and 0.51 (*beta0* and -*v* in loss task and reward task), indicating that all parameters contributed similarly. A higher value on this component reflected lower behavioural consistency. This component strongly correlated with both total score and choice cross-entropy, with a particularly stronger association with cross-entropy. This suggests that cross-entropy may better capture individual differences in decision-making processes, as it is less influenced by random fluctuations in early trials before learning stabilises, which can affect total scores. Among the brain state metrics, those that showed significant correlations with behavioural indices included the transition probabilities from states 2 to 1 and 3 to 2 (both negatively correlated), as well as the transition probability of remaining in state 4 (state 4 to 4) and the dwell time of state 4 (both positively correlated) (Supplementary Table. 5).

**Fig. 4.**
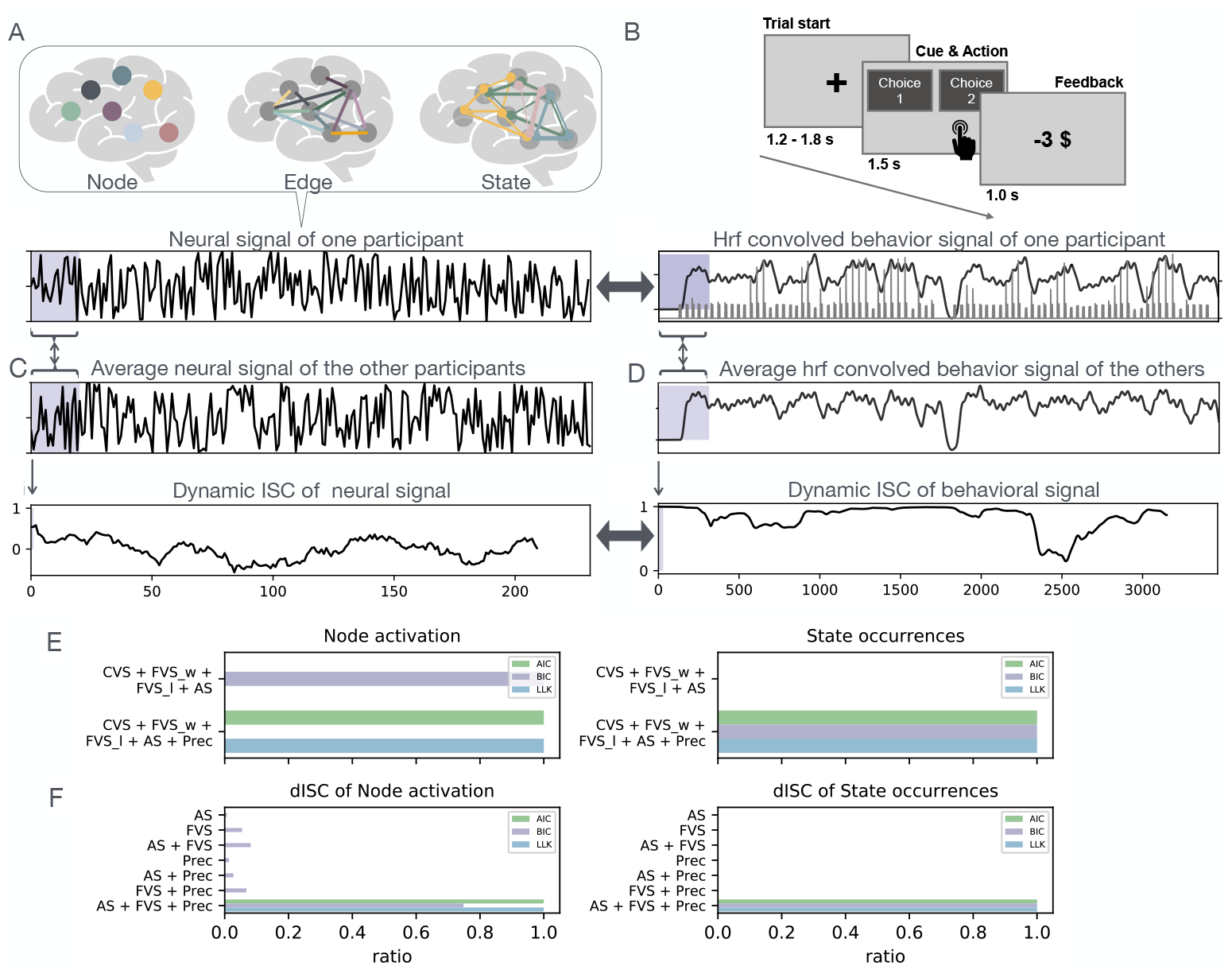
Schematic illustration of analysis for explaining neural signals by behavioural signals. (A) From functional magnetic resonance imaging (fMRI) data measured during decision-making tasks, neural signals are extracted at three levels. (B) The task begins with the gaze fixation on a central cross for a random duration ranging from 1.2 to 1.8 s; subsequently, the meaningful experimental stimulus is presented. The “Cue” refers to the presentation of two option figures and lasts for 1.5 s during which the participant had to choose one. After 1.5 s, the choice results are presented as the monetary reward or loss. We defined this presentation result as “Feedback” and lasts for 1.0 s. The images of each choice are shown as meaningless black-and-white pictures, and different sets of images are shown during each part of the task. Selection should be made in 1.5 s, during which the Cue images are presented on the screen. Policy precision signal is generated by the calculation process as in Figure 1D, then aligned to the time-point of Feedback presentation because the gradient update is assumed to occur by the free energy induced by Feedback. (C,D) By calculating the inter-subject correlation (ISC) of time-series over the sliding window, neural and behavioural dynamic inter-subject correlation (dISC) were obtained, respectively. (E) Performance comparison of conventional GLM. (F) Performance of dISC GLM. The x-axis represents the ratio of the number of regions where each model was measured as best (smallest values of each criterion), and the y-axis shows the independent variables included in each GLM model. CVS:cue visual stimulus; FVS:feedback visual stimulus; FVS w:feedback visual stimulus of win; FVS l:feedback visual stimulus of lose; AS:action selection; Prec:policy precision signal

**Fig. 5.**
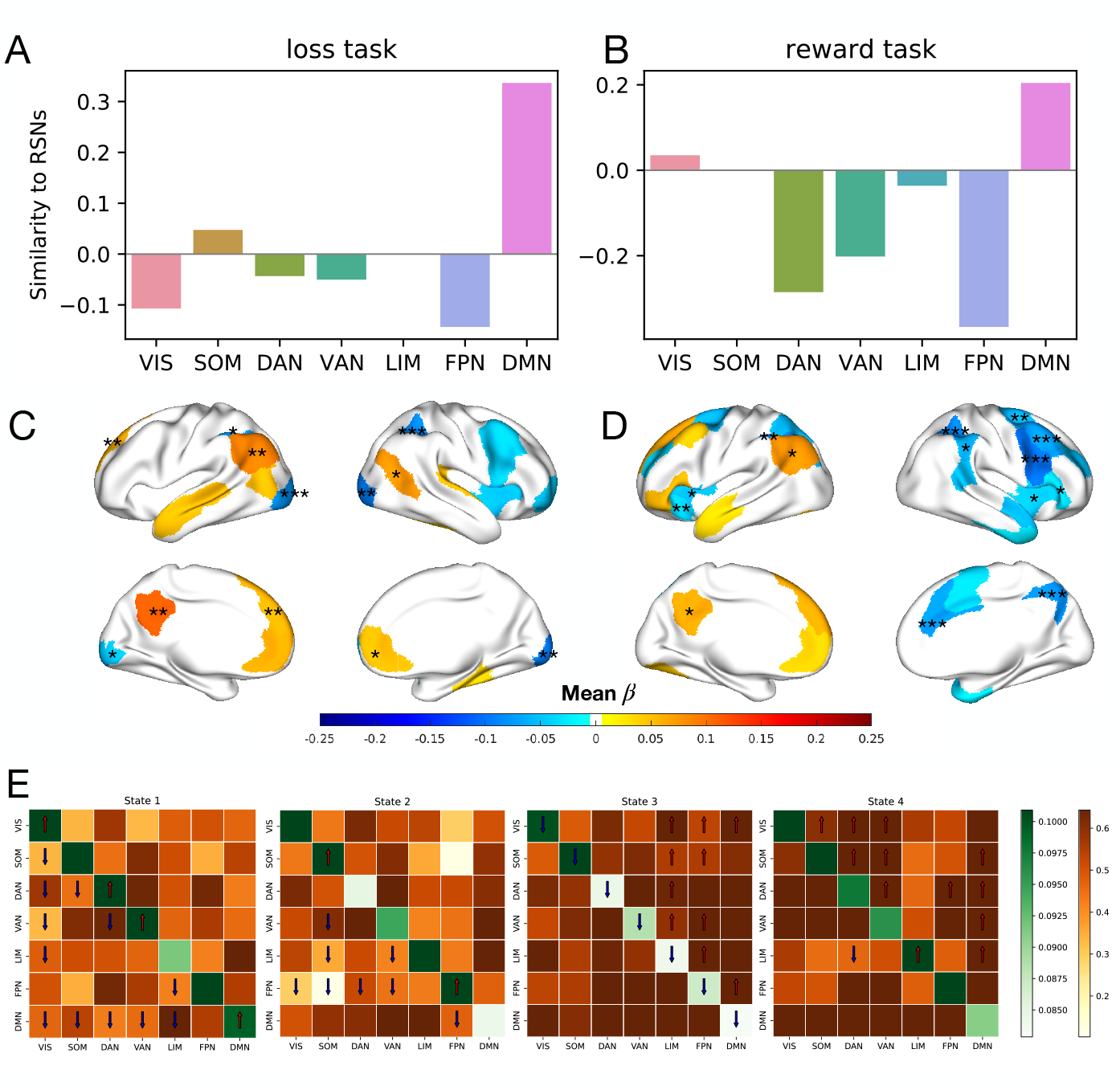
Results of a one-sample t-test using beta coefficient values result from GLM to investigate the regions related to the policy precision signal. (A, B) Cosine similarity between beta coefficient values of significantly related regions (p *<* 0.05) and the seven networks of intrinsic functional connectivity from [32]. Regions included within the default mode network (DMN) show positive correlations, while those included in the dorsal attention network (DAN) and frontoparietal network (FPN) show negative correlations. (C, D) Coloured regions have significant correlations with policy precision signal (p *<* 0.05). * indicate statistically significant predictors (p *<* 0.01), **, *** indicate remained significant after controlling for the false discovery rate (FDR) (** for q *<* 0.05, *** for q *<* 0.01). (E) Mean intra- and inter-connectivity between seven networks of each state calculated using adjacency matrices obtained from the Graph Laplacian mixture model. Upward arrows marked in the upper triangles of matrices indicate that those connectivities were highest among the states, whereas downward arrows marked in the lower triangles of matrices indicate that those were the lowest values among the states.

### 2.4 Model validation in population, including the patients with MDD

Finally, we determined whether the parameters estimated from the new model could better discriminate clinical characteristics. To this end, we conducted the behavioural task with a population that included patients diagnosed with MDD. The task composition was slightly modified by setting the probability of winning on the better bandit to 0.75 and mixing loss and reward tasks within the same session. Two task sets, 1 and 2, were used (n = 74 and n = 43, respectively). Model comparison among learning model families indicated that the beta distribution with a fixed decay parameter provided the best fit overall, except for the loss task in the task set 2 group (Supplementary Figure. 3). However, in that case, the difference between the best model and the second-best model (beta distribution with fixed decay) was not substantial. Therefore, for consistency, we adopted the beta distribution with a fixed decay model for all subsequent analyses.

We then compared action models under this learning model. Consistent with previous findings, the softmax model was the best-fitting model. However, in analyses that assessed how well parameter differences predicted CE, the softmax model performed poorly (Supplementary Figure. 4). This discrepancy became even more pronounced in this Experiment, which included patients, suggesting that greater inter-subject heterogeneity amplifies the limitations of the softmax model in explaining behavioural variability.

We conducted three analyses using the estimated behaviour parameter values from each model: (1) logistic regression predicting group membership between healthy controls (HC; n = 59) and patients with major depressive disorder (MDD; n = 58); logistic regression distinguishing non-suicidal ideation (NSI; n = 23) from suicidal ideation (SI; n = 35); (3) analysis of covariance (ANCOVA) predicting Hamilton Depression Rating Scale (HAM-D) scores [34] from the behavioural parameters. All analyses controlled for age, sex, education level, and task set. Behavioural parameters estimated from the loss and reward tasks of each model were initially included as independent variables. Due to high correlations, especially involving *β*_0_ (Supplementary Figure. 5A), we calculated the variance inflation factor (VIF) and iteratively excluded variables with VIF values exceeding 10 to mitigate multicollinearity. As a result, *β*_0_ was excluded from the EFE models, while both *w* and *β*_0_ were excluded from the WSLS-RL model, leaving four parameters in each model for final analysis. Parameters *util, ls*, and *β*_0_ from the reward task were significant predictors distinguishing between the control and MDD groups. In contrast, when differentiating between NSI and SI groups, significant predictors emerged from the loss task, and this pattern was observed only in the *efegc* model. Regarding the prediction of HAM-D scores, the results are similar to those of the HC vs. MDD logistic regression, but parameters only from the *efegc* and WSLS-RL models were significant. These findings suggest that these models may better capture individual-specific characteristics relevant to depressive symptoms.

## 3 Discussion

To explain psychiatric symptoms or real human cognitive actions, it is essential to utilise biologically plausible models rather than relying solely on those that demonstrate the best performance. This approach ensures that the model is accurate and reflective of the complexities of neural processes. Our model included distributional representation during the learning process, calculated expected energy during action selection, and optimised policy precision by observing each trial. Although this may seem overly complex, including biologically plausible parameters and signals significantly enhances explanatory power in neural signal data and validates the advantage in the clinical data.

The main advantage of this model lies in its ability to account for the context-dependent interplay between MF and MB strategies using a single free energy function, thereby avoiding additional weighting parameters. By optimising policy precision based on outcomes from previous trials, the model not only improved task performance but also enhanced the explanatory power of the free parameter in capturing clinical characteristics (Figure 6). Furthermore, the rate of change in policy precision emerged as a meaningful signal in explaining neural dynamics (Figure 4E), particularly when inter-subject correlations (ISC) were considered, leading to improved explanatory power regarding variability in neural responses (Figure 4F).

**Fig. 6.**
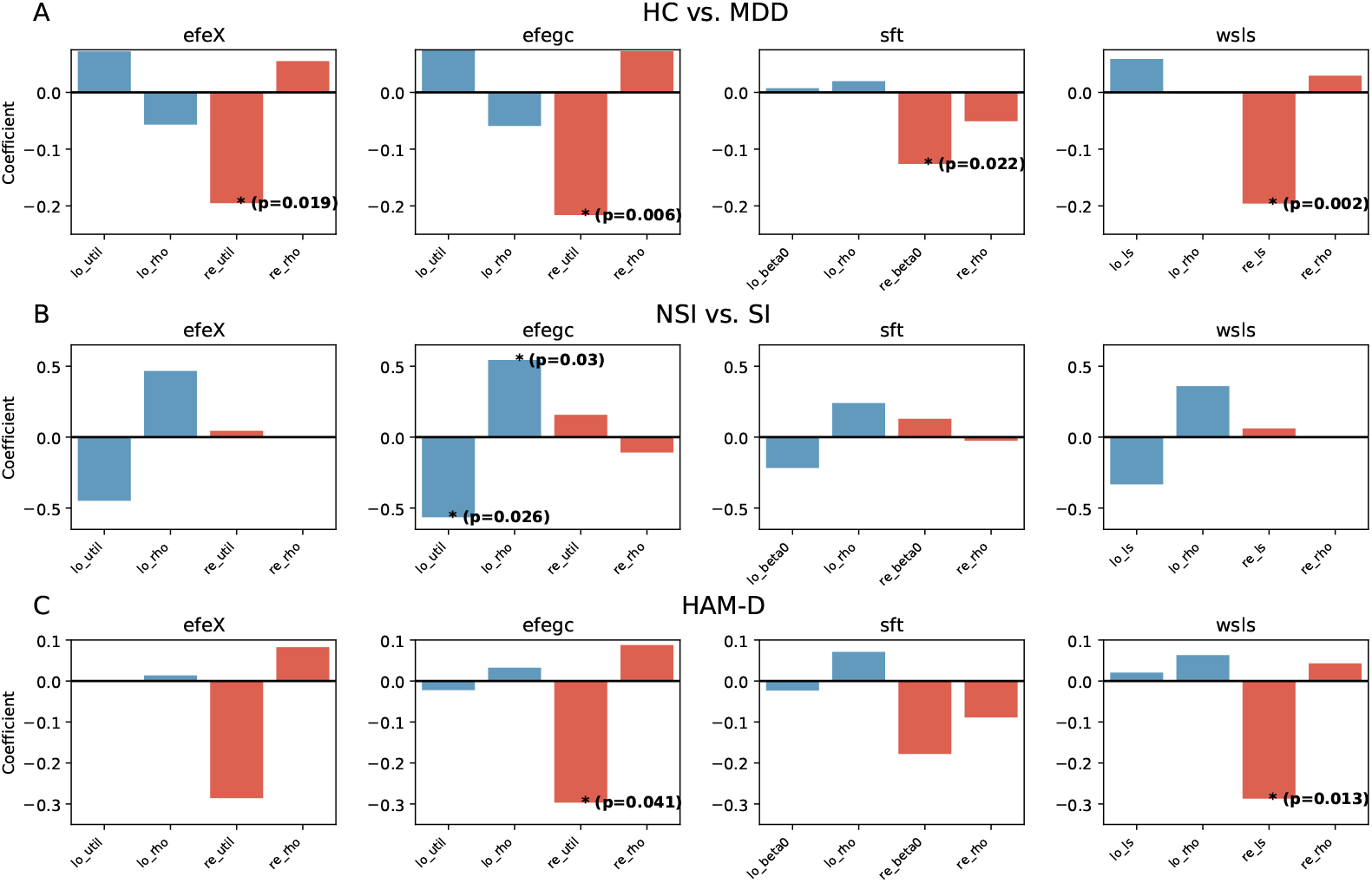
Results of three types of analyses using parameter values estimated from four behaviour models. (A, B) Results of logistic regression analysis predicting group using parameters. (C) Results of analysis of covariance (ANCOVA) predicting HAM-D scores and each individual parameter value, respectively. Asterisks (*) indicate statistically significant predictors (p *<* 0.05).

The policy precision signal plays a critical role in the decision-making process by modulating the weight of MB-dependent terms. In the AIF, policy selection is assumed to be a fully model-based process [35]. From this theoretical standpoint, equating the F term with MF control is not accurate. However, we adopted this interpretation since MF control may be conceptualised as a limiting case on the MB–MF continuum—one in which the influence of an internal model is substantially reduced. Similarly, it may not be entirely precise to equate the WSLS strategy with MF control in the WSLS-RL model. Although the commonly used MB–MF dichotomy associates MB strategies with goal-directed behaviour and MF strategies with habitual behaviour (a simplification often used for intuitive clarity), this distinction is not entirely accurate, and MF strategies can be considered goal-directed [36–39]. Nevertheless, for consistency with the conventional interpretation, we refer to WSLS and related strategies as “MF”. Additionally, in AIF, habitual behaviour is represented separately as an additional term (often referred to as the E term). In the context of our task, this would correspond to increasing the probability of selecting an alternative option following an incorrect choice, effectively altering the model such that the free energy of the previously selected (incorrect) option increases. This mechanism is already captured in our formulation (Equation 11); therefore, we opted not to explicitly include the habitual component, as it would substantially overlap with the F term and is unlikely to provide meaningful improvements in model performance.

In the neural signal analysis, regions with a positive correlation with the policy precision signal were predominantly affiliated with the DMN, whereas those with a negative correlation were associated with the DAN and the FPN. These regional findings align with network-level results, which revealed a positive correlation with the occurrence of State 2 (dominated by the DMN) and a negative correlation with State 3 (dominated by the DAN and FPN). The anti-correlation between these states has been consistently observed in prior studies, where the DMN is typically associated with the resting (i.e., task-negative) state, and the DAN and FPN with task-positive processing. Although the DMN is generally activated during rest, mind-wandering, and self-referential thought [40–42], recent evidence suggests its role extends to the processing of intrinsic information during task performance [43–46]. In line with this, State 2, dominated by DMN activity, was the second most frequently observed state during task execution in our study Supplementary Figure. 12B, supporting the functional involvement of the DMN during goal-directed behaviour. The DMN has been suggested to facilitate the detection of associative relevance between internal and external stimuli and contributes to value coding [47]. From this perspective, an alternative proposal argues that the DMN supports recursive exploration by selecting optimal actions within environmental contexts, potentially operating at higher levels of the inferential hierarchy to minimise free energy at lower levels [48, 49]. Additionally, the hippocampus, which was active in State 2 (Supplementary Figure. 1J), is implicated in internal model construction and planning [50, 51], further linking DMN functionality with internal modelling processes. Conversely, activation of State 3, characterised by dominant DAN and FPN activity, was associated with decreased policy precision—likely reflecting increased uncertainty and the need for externally directed attention.

Correlation analyses between behavioural parameters inferred from the model and state dynamic metrics revealed that longer dwell time in State 4 was associated with increased choice inconsistency. In contrast, transition probabilities related to State 2 showed the opposite pattern. Further examination revealed a mutually inhibitory relationship between States 2 and 4: an increase in State 2 dwell time was associated with reduced transitions involving State 4, and vice versa (Supplementary Figure. 12D). This suggests that while anti-correlation between States 3 and 2 related to the policy precision signal may reflect a cooperative dynamic supporting task performance via policy precision modulation, State 4 may interrupt the functional role of State 2. State 4, despite its generally lower nodal activation, showed relatively higher activation in the LIM and VAN and was characterised by high connectivity (Figure 5E). Previous studies suggest that VAN is activated by surprising external stimuli [52, 53] and plays a role in regulating the balance between the DMN and DAN [54, 55]. Conversely, DAN activity has been shown to suppress VAN, supporting sustained goal-directed attention. The stronger this suppression, the better the task performance [56]. Based on this, we speculate that when surprise-induced VAN activation occurs due to unexpected outcomes, a rapid suppression by DAN may be necessary to restore attentional focus and maintain learning based on the internal model. This dynamic may reduce the dwell time in State 4 and facilitate appropriate engagement of States 2 and 3, thereby improving task performance and behavioural consistency. Moreover, these results align with the view that MF strategies are not a distinct category opposed to MB control, but rather reflect a relative state over the MB continuum, where MB influence is diminished. From this perspective, transitions between States 2 and 3 correspond to a shift in action strategy characterised by reduced MB control and increased reliance on alternative mechanisms. Importantly, ‘MF reliance’ does not necessarily imply that participants actively engage in a distinct MF strategy; rather, it indicates a state in which MB reliance is weakened. The engagement of task-positive regions observed in State 3 may serve a compensatory role, supporting decision-making in the face of reduced MB influence.

In a related study by [57], a single latent dimension termed “decision acuity” was identified, capturing shared variance across multiple decision-making tasks. This construct aligns closely with the prior policy precision (*beta0*) or its principal component in the active inference framework. Their findings showed that intra- and inter-connectivity within FPN, medial prefrontal cortex (MPC), orbitofrontal cortex, medial and lateral (OFC), opercular cortex (OPC) posterior cingulate cortex (PCC) right dorsolateral prefrontal cortex (RDC) somatosensory and motor areas (SMT) and visual regions modules significantly predicted decision acuity. As shown in Figure 5E, State 4 exhibited high integration within RSNs overlapping these modules. Although these results are from resting-state fMRI, such baseline connectivity patterns likely influence task-state neural dynamics, suggesting a degree of relevance to our findings.

Moreover, while task-fMRI was not conducted in Experiment 2, the observed behavioural parameter differences between healthy controls and individuals with MDD or SI-MDD may be attributable to differences in brain state dynamics. Specifically, modulating State 4 dwell time to maintain levels similar to those observed in healthy individuals during surprise events may aid in symptom alleviation or serve as an objective marker for assessing MDD risk.

The parameters *beta0* and *util* in reward decision were significantly reduced in patients with MDD. These findings are consistent with previous studies reporting impairments in the utilization of reward value in patients with MDD [58, 59]. In contrast, the corresponding parameters from the loss domain appeared relatively preserved when compared to healthy controls. However, they emerged as significant predictors of suicidal ideation within the MDD group, and this pattern was observed exclusively in the *efegc* model (*β*=−1.412, *p*=0.026 for *util, β*=1.36, *p*=0.030 for *ρ*). This additional reduction in loss aversiveness (*util* in the loss task) may reflect an increased propensity towards self-harm or suicidal ideation, suggesting that altered sensitivity to negative outcomes could be key features in identifying suicide risk among individuals with MDD. However, since this subgroup analysis included a small number of people for each NSI and SI group (n = 23 and 35, respectively), future studies with larger samples remain warranted to confirm our findings.

A behaviour model that can reflect the various mental states of people is warranted to better explain mental states through the model. In addition, there are limitations to understanding individual characteristics through parameters obtained from behavioural modelling. Therefore, high-dimensional brain signal data that reflects individual diversity must be integrated. This makes behavioural modelling even more critical. Rather than focusing solely on better explaining behavioural outcomes, we must prioritise the development of biologically plausible models, identify their neural correlates through brain signals, and understand mental symptoms in terms of the differences or abnormalities that arise from these connections. This will explain the differences in various brain states through models and serve as a foundation for personalised neuromodulatory treatment through accurate correlation between behaviour and individual brain activity.

## 4 Materials and methods

### 4.1 Participants

#### 4.1.1 Experiment 1

A total of 54 volunteers participated in Experiment 1 (15 females and 39 males; mean [standard deviation] age: 22.4 [3.2] years; all participants were right-handed and had no history of psychiatric or neurological disorders). All participants provided written informed consent for all the procedures and data usage before the start of the study. The experimental procedures were approved by the Ethics Committee of Korea Advanced Institute of Science and Technology.

#### 4.1.2 Experiment 2

A total of 58 patients with MDD (35 SI and 23 NSI) and 59 HCs without any current or history of psychiatric illness, aged 18 to 34 years, were recruited through the outpatient clinic of the Samsung Medical Center in Seoul, South Korea, between July 2018 and October 2020. MDD was diagnosed by two psychiatrists (H. Kim and H.J. Jeon) based on clinical interviews. Those whose diagnosis was categorised as MDD per DSM-5 were included in the MDD group. The exclusion criteria were: (1) MDD with psychotic features; (2) comorbidity of any other psychiatric illnesses including bipolar disorder, schizophrenia, delusional disorder, delirium, neurocognitive disorder, intellectual disability, and other mental disorders due to another medical condition;

history of substance-related disorders except for tobacco-related disorders within 12 months; (4) primary neurologic illness or history of brain damage; (5) a history of major physical illness. HCs were recruited from the community using an advertisement for the Clinical Trial Center of Samsung Medical Center. The control group enrolled those who had no history of major depressive episodes and scored ≤ 7 on the HAM-D scale, using the same exclusion criteria that were applied to participants in the MDD group. The study design was approved by the Institutional Review Board of Samsung Medical Center (IRB No. 2018-04-137). All participants provided written informed consent before participating in this study in accordance with the Declaration of Helsinki. SI was assessed with the suicidality module of the Mini International Neuropsychiatric Interview [60]. Among the patients with MDD, the SI group included participants who answered “yes” to the question “Did you think about suicide during the past one month?” (SI group, n=35), whereas the no SI group included those who answered “no” to the same question (NSI group, n=23). Depression severity was measured using the 17 items in HAM-D [34].

### 4.2 fMRI data acquisition

In Experiment 1, all participants underwent MRI using 3.0-T MRI (Siemens Verio Syngo Scanner). Both T1-weighted (T1w; echo time [TE] = 2.02 ms, repetition time [TR] = 2,400 ms, field of view (FOV) = 224 × 224 mm, and 0.7 mm iso-voxel) and T2-weighted (T2w; TE = 330 ms, TR = 2200 ms, FOV = 224 × 224 mm, and 0.7 mm iso-voxel) structural images were scanned. A total of 240 echo-planar imaging scans of BOLD responses (multiband factor = 4, TE = 32 ms, TR = 1.5 s, flip angle = 50°, FOV = 225 × 221 mm, matrix size = 110 × 107, 2.0 mm iso-voxel with no gap, 64 slices, phase encoding = anterior to posterior) were performed using a 32-channel head coil. Field map images (TR = 731 ms, TE: 4.92 ms and 7.38 ms, flip angle = 90°) were also acquired to correct distorted images. In every trial, the participants performed the experimental task with both hands to press an MRI-compatible button to make their decision. In the pretraining session, prior to the first scan, the participants were acquainted with the apparatus and tasks performed during scanning. Sequences executed during this pretraining session were not encountered later in the experiment. Preprocessing and processing of the neuroimaging data were performed using AFNI (v23.1.00) [61, 62], FSL (v6.0.5.2), FreeSurfer (v7.1.1), Workbench (v1.5.0), and Advanced Normalization Tools (ANTs). Following the Human Connectome Project, a minimal processing pipeline for structural images, structural T1w and T2w images [63], were processed, involving automatic segmentation of surfaces and reconstruction using FreeSurfer 7.1.1 (https://surfer.nmr.mgh.harvard.edu/) [64, 65]. The generated volumes and surface files were converted to the CIFTI file format using ciftify (https://edickie.github.io/ciftify/) [66] including MSMSulc surface realignment [67].

Raw time-series of task fMRI were performed with slice-timing correction, realignment, and skull stripping using AFNI commands (3dTshift, 3dvolreg, and 3dAutomask) [68]. Realigned data were unwarped using a fieldmap in FSL. A bandpass filter (highpass = 0.01) was applied to the unwarped functional data and motion artifact time-series before regression. Using algorithms in ANTs, we registered the unwarped functional data into a 2-mm MNI152 T1w template space via skull-stripped T1w images pre-processed with AFNI. The eroded white matter (WM) and ventricular cerebrospinal fluid masks segmented using FreeSurfer were moved to the functional image space using the transformation matrix generated via ANTs in the previous step. PCA (3dpca in AFNI) was applied to the eroded ventricular mask in bandpass-filtered, scaled functional data, and ventricular PCA time-series was acquired. Using the fast ANATICOR algorithm in AFNI [69] on an eroded WM mask, a voxel-wise local WM regressor was generated.

3dTproject in AFNI, all regressors generated were projected into a regression matrix. Finally, the bandpass-filtered, scaled functional data were denoised, including pre-whitening, to remove the autocorrelation property using 3dREMLfit (residual maximum likelihood) in AFNI. Using transformation matrices and warping images generated by ANT algorithms, the denoised functional data were directly moved to a 2-mm MNI152 T1w template space and projected to the CIFTI format using the ciftify package [66].

### 4.3 Neural signal extraction

After preprocessing, we acquired the averaged ROI time-series extracted from the Schaefer Atlas [70], including 100 regions covering the cortex, and the Melbourne Subcortex Atlas [71] covering the subcortex. Additionally, to include other subcortical regions related to the release of important neurotransmitters, including dopamine and norepinephrine, we added some regions to the CIT168 atlas [72] and Harvard Ascending Arousal Network Atlas [73] to obtain signals from a total of 147 ROIs.

To extract the representation of overall brain states associated with network-level interactions, we employed the Graph Laplacian mixture model (GLMM). This method identifies meta-networks representing both intra- and inter-network interactions by estimating the Laplacian matrices that characterise the graph structure and dynamics of brain states. This method showed state classifications consistent with the timing of various task paradigms in the HCP dataset [31, 74]. GLMM provides information on the brain state activation patterns (estimating means), functional connectivity structure (Laplacians), and their temporal dynamics (probability of occurrences over time). Four major states were identified and depicted in Supplementary Figure. 1.

### 4.4 Behaviour task and modelling

#### 4.4.1 Experiment 1

Participants faced two choices: one with a 70% likelihood of winning and the other with a 30%. The number of trials was 40 per part; thus, a total of 80 trials were performed by each participant. Participants were explicitly instructed that one of the two bandits had a higher probability of winning, and were asked to infer which one was better to win as the trial progressed to maximise the accumulated reward. Participants were expected to learn about the reward value of each option by trial-and-error and utilise computed reward values to make their decision. Therefore, the probability of one choice being better represents the hidden state value inferred by the agents during the task. This state could be modelled using either a beta or a normal distribution. The inferred value over one choice could be updated in every trial, even if that choice was not selected, because winning or losing by selecting one choice simultaneously means losing or winning by selecting the other choice. The expected value can be represented by either scalar probability or a distribution form. Using a scalar value corresponds to the Rescorla-Wagner model. Meanwhile, using a distribution form, we can utilise the variance of state when calculating behavioural measures in the model, such as learning rate or free energy. Each trial corresponds to Bernoulli trial with outcomes of win or lose, and state value *q*(*θ*) can be represented by a beta distribution, where *θ* is the probability of one choice being better bandit. The beta distribution is parameterised as:

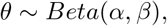

where *α* and *β* are equal to one plus the number of win or lose outcomes, respectively, for a given choice. In this case, the learning process as the task progresses can be represented simply by adding one according to the outcome of each trial (*o*_*t*_), as the likelihood of outcome at each trial is a binomial distribution, which is conjugate to the beta distribution:

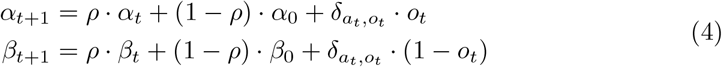

*δ* indicates the Kronecker delta: 1 if both variables *a*_*t*_ and *o*_*t*_ are equal (when selecting choice *1* results in a win or selecting choice *0* results in a lose), and 0 otherwise. *ρ* is the decay parameter (0 *< ρ <* 1). Smaller values of *ρ* lead the concentration parameters to converge toward their initial values, effectively corresponding to a forgetting process in which learning from observations fades over time. This parameter can be dynamically updated across trials as follows:

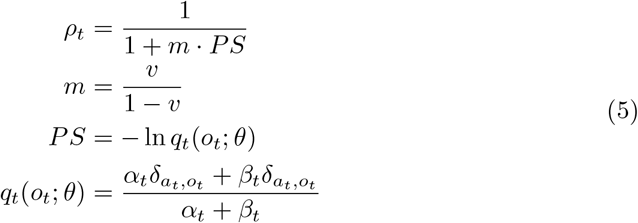

The likelihood of observing *o*_*t*_ given the current belief is denoted as *q*_*t*_(*o*_*t*_; *θ*), which can be efficiently computed under a beta distribution. When an outcome *o*_*t*_ that was unlikely under the current belief is observed, the level of surprise increases. This mismatch between prior beliefs and observed outcomes is quantified as predictive surprise (PS). An increased PS reduces the relative influence of the prior belief during belief updating, thereby allowing the new observation to exert a stronger effect on the posterior belief. Meanwhile, the parameter *m*, which depends on the volatility parameter (0 *< v <* 1), modulates how strongly the surprise influences learning. Volatility refers to the degree to which the underlying generative probability has drifted, making it less compatible with the current belief. Higher volatility implies greater environmental instability, and thus a stronger need to adapt by giving more weight to recent observations.

The state value *q*(*θ*) can be represented by a normal distribution as *θ* ~ *Norm*(*µ, σ*^2^) with a prior mean of 0.5 and typically a large standard deviation of 10. In this case, corresponding to the Kalman-filter model [26], likelihood of outcome at each trial can be represented as normal distribution, and the standard deviation of the observation (*σ*_*o*_ in Equation 6) either can be fixed or defined as free parameter. That modulates how much prior belief will be updated:

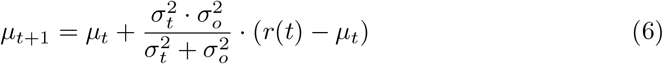

As an action model, we used a conventional softmax function (Equation 2), WSLS-RL (Equation 3), and two models based on AIF scheme utilising EFE. Formally, we express the EFE of a choice *a* on trial *t* as:

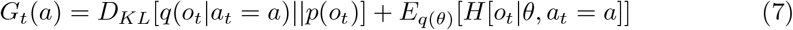

preference distribution of outcomes, *p*(*o*_*t*_), is encoded using *util* parameter for each participant as

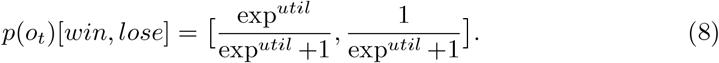

The probability of selecting choice *a* is then calculated as:

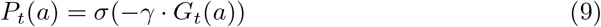

In addition, the precision of beliefs about actions *γ*, itself can be inferred instead of a fixed value [75]. In the *efeX* model, the following adjustment excludes the *beta0* parameter (*β*_0_) which is used as *γ* in Equation 9, whereas in the *efegc* model, the initial value of *γ* is calculated from the fixed prior value of *β*_0_ [76]. Thus, the probability that the agent will choose each policy *π*_0_ will be

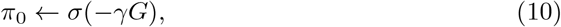

As precision weights the EFE to determine the probability of choosing each policy, the results of trial *t* are given as either a win or a loss, and the variational free energy (VFE) of each policy *F* over that observation is calculated. Subsequently, the probability of choosing each policy after a subsequent observation is:

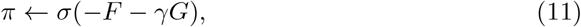

and the error of the agent’s EFE (*G*_*error*_) is given as:

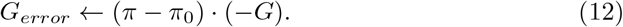

The above equation reflects the level of (dis)agreement between the EFE (*G* and *π*_0_) and the VFE of the observation (*F* and *π*) [76]. Thus, the *β* whose inverse means *γ* is updated based on *G*_*error*_.

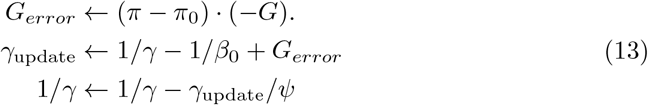

Therefore, policy precision reflects the reliability of the EFE based on new observations that represent either winning or losing outcomes. If the result of a chosen action betrays expectations, the precision decreases, causing the probability of action selection to rely less on the expected free energy. Simultaneously, this increases the effect of free energy that is calculated from the previous outcome. This process explains the interplay between model-based and model-free strategies.

#### 4.4.2 Experiment 2

The overall paradigm of the decision-making task is similar to that of Experiment 1. However, participants performed 80 trials at once, and each trial’s decision was either a loss or reward specific. In reward decision-making task trials, choice options are presented with a blue boundary, while red boundaries are used in loss trials (Supplementary Figure 2). One option was associated with a high probability of reward (75% of reward) while another option had a high probability of non-reward. Reward and loss schedules were semi-randomised and counterbalanced with two schedules (74 participants for set 1 and 43 for set 2). The same behaviour models were used as in Experiment 1, with the addition of one learning model that considered decay along the trial sequence. For example, the second and third loss tasks occurred on the third and ninth trials of the 80-trial sequence. Considering decaying as the number of trials between those two trials, it is as many as six trials instead of one. Therefore,

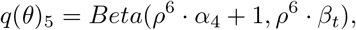

when selecting choice *a* results in a win.

### 4.5 GLM analysis

To identify the shared dynamics of neural engagement across participants who performed the same experimental task, we calculated the dISC for each level of the time-series [21, 77]. A sliding window with a length of three repetition times (TRs) corresponded to approximately the length of one trial (3.7-4.3s) and a 1 TR overlap. This was calculated by correlating the neural signal of one participant for each region within each window with the mean signal within the same window for the same region from the remaining participants (leave-one-out method) (Figure 4C). The ISC method increases the signal-to-noise ratio (SNR) to detect stimulus-induced inter-regional correlations. The improvement in the SNR arises from filtering out intrinsic neural dynamics (for example, responses arising from intrinsic cognitive processes unrelated to ongoing stimulus processing) and non-neural artifacts (for example, respiratory rate and motion) that can influence network correlation patterns within the brain but are not correlated across participants. The time-series of neural signal from node i of participant j is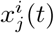; this signal consists of a linear combination of the shared component c(t), idiosyncratic responses to the stimulus for participant j 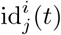, and spontaneous activity unrelated to the stimulus 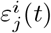. Therefore, the time series of participants a and b can be expressed as Equation [14], with shared c(t).

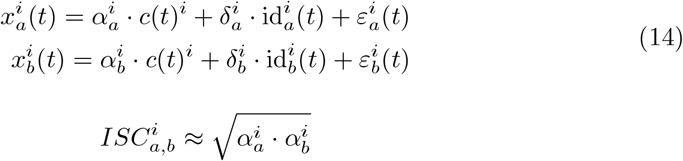

In the previous ISC study, the correlation between participants a and b was approximated to the 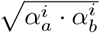 because id(t) and *ε*(*t*) are not systematically correlated across participants, whereas c(t) is perfectly correlated [21]. However, in the current study, the idiosyncratic response conforms to the policy precision signal, in that id(t) is not perfectly uncorrelated across participants. This is shown in the bottom plot of Figure 4D showing the ISC of the policy precision signals varied from 0 to 1, with a considerably high value. In the original logic of ISC, this idiosyncratic response should be averaged to a small value close to zero due to its inconsistent characteristic, such that only c(t) can be isolated; however, correlations across idiosyncratic responses of the participants exist. Therefore, the neural ISC values corresponded to the values including not only the effect of the shared component but also the idiosyncratic response component in the extent of ISC of the policy precision signal for participant *j*. Thus, the neural ISC value of each node *i* can be approximated as:

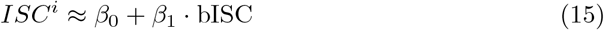

where we can assume that the intercept term *β*_0_ corresponds to the effect of *α*^*i*^ in Equation [14] and the *β*_1_ corresponds to the effect of ISC of behavioural signal (bISC), such as policy precision. If the bISC is 0, the result is equivalent to the third line of Equation [14]. This can be applied to the time-series data of each participant (*j*) using dynamic ISC (dISC) of neural and dISC of behavioural signals (dbISC in Equation [16]) for performing GLM analysis.

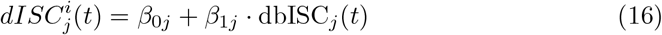

Since action timings (AS) and feedback visual stimulus (FVS) are also different across participants, we performed a GLM analysis that included dISC of these signals, both with and without dISC of policy precision signal. We then compared the model performance using AIC, BIS, and deviance scores.

To validate the relevance of policy precision signals in explaining neural signals, we performed a conventional GLM to identify which regions or interactions are related to this signal. Using beta coefficient values estimated from the GLM, we conducted a one-sample t-test to examine if those values are significantly positive or negative.

## 5 Conclusion

Our findings demonstrate that the biologically plausible AIF model provides a better reflection of individual states than conventional RL models. This highlights the potential of developing behavioural models in this manner to advance precision medicine in psychiatry.

## 6 Supplementary information

[Supplementary Materials.pdf](https://docs.google.com/uc?export=download&id=1W0wDOI4NbyvFxe5W1JJSFi2GyCwq_YIC)

## 8 Acknowledgements

This study was supported by the Medical Scientist Training Program from the Ministry of Science & ICT of Korea, and the Brain Research Program through the National Research Foundation of Korea (NRF) funded by the Ministry of Science & ICT (NRF-2022M3E5E8081200 and NRF-2016M3C7A1914448). We would like to thank Editage (www.editage.co.kr) for English language editing.

## 9 Data Availability

The data underlying this article will be shared upon reasonable request to the corresponding author. Codes used in the current study soon will be available at https://github.com/ydy-dyd-y/AIFdFC

## References

[1] Montague, P.R., Dolan, R.J., Friston, K.J., Dayan, P.: Computational psychiatry (2012). 10.1016/j.tics.2011.11.018

[2] Huys, Q.J.M., Maia, T.V., Frank, M.J.: Computational psychiatry as a bridge from neuroscience to clinical applications. Nature Publishing Group (2016). 10.1038/nn.4238

[3] Chen, C., Takahashi, T., Nakagawa, S., Inoue, T., Kusumi, I.: Reinforcement learning in depression: A review of computational research. Elsevier Ltd (2015). 10.1016/j.neubiorev.2015.05.005

[4] Maia, T.V., Frank, M.J.: From reinforcement learning models to psychiatric and neurological disorders. In: Nature Neuroscience, vol. 14 (2011). 10.1038/nn.2723

[5] Ahn, W.Y., Busemeyer, J.R., Wagenmakers, E.J., Stout, J.C.: Comparison of decision learning models using the generalization criterion method. In: Cognitive Science, vol. 32 (2008). 10.1080/03640210802352992

[6] Worthy, D.A., Hawthorne, M.J., Otto, A.R.: Heterogeneity of strategy use in the Iowa gambling task: A comparison of win-stay/lose-shift and reinforcement learning models. Psychonomic Bulletin and Review 20(2) (2013) 10.3758/s13423-012-0324-9

[7] Ahn, W.Y., Vasilev, G., Lee, S.H., Busemeyer, J.R., Kruschke, J.K., Bechara, A., Vassileva, J.: Decision-making in stimulant and opiate addicts in protracted abstinence: Evidence from computational modeling with pure users. Frontiers in Psychology 5(AUG) (2014) 10.3389/fpsyg.2014.00849

[8] Worthy, D.A., Todd Maddox, W.: A comparison model of reinforcement-learning and win-stay-lose-shift decision-making processes: A tribute to W.K. Estes. Journal of Mathematical Psychology 59(1) (2014) 10.1016/j.jmp.2013.10.001

[9] Yechiam, E., Busemeyer, J.R., Stout, J.C., Bechara, A.: Using cognitive models to map relations between neuropsychological disorders and human decisionmaking deficits. Psychological Science 16(12) (2005) 10.1111/j.1467-9280.2005.01646.x

[10] Dombrovski, A.Y., Szanto, K., Clark, L., Reynolds, C.F., Siegle, G.J.: Reward signals, attempted suicide, and impulsivity in late-life depression. JAMA Psychiatry 70(10) (2013) 10.1001/jamapsychiatry.2013.75

[11] Cohen, J.D., McClure, S.M., Yu, A.J.: Should I stay or should I go? How the human brain manages the trade-off between exploitation and exploration. In: Philosophical Transactions of the Royal Society B: Biological Sciences, vol. 362 (2007). 10.1098/rstb.2007.2098

[12] Schulz, E., Gershman, S.J.: The algorithmic architecture of exploration in the human brain (2019). 10.1016/j.conb.2018.11.003

[13] Parr, T., Pezzulo, G., Friston, K.J.: Active Inference: the Free Energy Principle in Mind, Brain, and Behavior. MIT Press, ??? (2022)

[14] Rescorla, R.A., Wagner, A.R.: A Theory of Pavlovian Conditioning: Variations in the Effectiveness of Reinforcement and Nonreinforcement. Clasical conditioning II: current research and theory (1972)

[15] Findling, C., Skvortsova, V., Dromnelle, R., Palminteri, S., Wyart, V.: Computational noise in reward-guided learning drives behavioral variability in volatile environments. Nature Neuroscience 22(12) (2019) 10.1038/s41593-019-0518-9

[16] Herrnstein, R.J., Rachlin, H., Laibson, D.I.: The matching law: Papers in psychology and economics. New York: Cambridge, MA. Kacelnik, A 53 (1997)

[17] Otto, A.R., Taylor, E.G., Markman, A.B.: There are at least two kinds of probability matching: Evidence from a secondary task. Cognition 118(2) (2011) 10.1016/j.cognition.2010.11.009

[18] Steyvers, M., Lee, M.D., Wagenmakers, E.J.: A Bayesian analysis of human decision-making on bandit problems. Journal of Mathematical Psychology 53(3) (2009) 10.1016/j.jmp.2008.11.002

[19] Kim, D., Park, G.Y., ODoherty, J.P., Lee, S.W.: Task complexity interacts with state-space uncertainty in the arbitration between model-based and modelfree learning. Nature Communications 10(1) (2019) 10.1038/s41467-019-13632-1

[20] Kraskov, A., Stögbauer, H., Grassberger, P.: Estimating mutual information. Physical Review E - Statistical Physics, Plasmas, Fluids, and Related Interdisciplinary Topics 69(6) (2004) 10.1103/PhysRevE.69.066138

[21] Nastase, S.A., Gazzola, V., Hasson, U., Keysers, C.: Measuring shared responses across subjects using intersubject correlation. Social Cognitive and Affective Neuroscience 14(6) (2019) 10.1093/scan/nsz037

[22] Turecki, G., Brent, D.A., Gunnell, D., O’Connor, R.C., Oquendo, M.A., Pirkis, J., Stanley, B.H.: Suicide and suicide risk (2019). 10.1038/s41572-019-0121-0

[23] Brown, V.M., Wilson, J., Hallquist, M.N., Szanto, K., Dombrovski, A.Y.: Ventromedial prefrontal value signals and functional connectivity during decision-making in suicidal behavior and impulsivity. Neuropsychopharmacology 45(6) (2020) 10.1038/s41386-020-0632-0

[24] Dombrovski, A.Y., Hallquist, M.N., Brown, V.M., Wilson, J., Szanto, K.: Value-Based Choice, Contingency Learning, and Suicidal Behavior in Mid- and Late-Life Depression. Biological Psychiatry 85(6) (2019) 10.1016/j.biopsych.2018.10.006

[25] Clark, L., Dombrovski, A.Y., Siegle, G.J., Butters, M.A., Shollenberger, C.L., Sahakian, B.J., Szanto, K.: Impairment in Risk-Sensitive Decision-Making in Older Suicide Attempters With Depression. Psychology and Aging 26(2) (2011) 10.1037/a0021646

[26] Kalman, R.E.: A new approach to linear filtering and prediction problems. Journal of Fluids Engineering, Transactions of the ASME 82(1) (1960) 10.1115/1.3662552

[27] Liakoni, V., Modirshanechi, A., Gerstner, W., Brea, J.: Learning in volatile environments with the bayes factor surprise. Neural Computation 33(2) (2021)

[28] Friston, K., Mattout, J., Trujillo-Barreto, N., Ashburner, J., Penny, W.: Variational free energy and the Laplace approximation. NeuroImage 34(1) (2007) 10.1016/j.neuroimage.2006.08.035

[29] Rigoux, L., Stephan, K.E., Friston, K.J., Daunizeau, J.: Bayesian model selection for group studies - Revisited. NeuroImage 84 (2014) 10.1016/j.neuroimage.2013.08.065

[30] Penny, W.D., Stephan, K.E., Daunizeau, J., Rosa, M.J., Friston, K.J., Schofield, T.M., Leff, A.P.: Comparing families of dynamic causal models. PLoS Computational Biology 6(3) (2010) 10.1371/journal.pcbi.1000709

[31] Ricchi, I., Tarun, A., Maretic, H.P., Frossard, P., Van De Ville, D.: Dynamics of functional network organization through graph mixture learning. NeuroImage 252 (2022) 10.1016/j.neuroimage.2022.119037

[32] Thomas Yeo, B.T., Krienen, F.M., Sepulcre, J., Sabuncu, M.R., Lashkari, D., Hollinshead, M., Roffman, J.L., Smoller, J.W., Zöllei, L., Polimeni, J.R., Fisch, B., Liu, H., Buckner, R.L.: The organization of the human cerebral cortex estimated by intrinsic functional connectivity. Journal of Neurophysiology 106(3), 1125– 1165 (2011) 10.1152/jn.00338.2011

[33] Vidaurre, D., Abeysuriya, R., Becker, R., Quinn, A.J., Alfaro-Almagro, F., Smith, S.M., Woolrich, M.W.: Discovering dynamic brain networks from big data in rest and task (2018). 10.1016/j.neuroimage.2017.06.077

[34] Yi, J.S., Bae, S.O., Ahn, Y.M., Park, D.-b.: Validity and Reliability of the Korean Version of the Hamilton Depression Rating Scale (K-HDRS). Journal of the Korean Neuropsychiatric Association 44(4) (2005)

[35] Thomas Parr, Giovanni Pezzulo, Karl J. Friston: Active Inference. Technical report (2022)

[36] Friston, K., FitzGerald, T., Rigoli, F., Schwartenbeck, P., O’Doherty, J., Pezzulo, G.: Active inference and learning. Elsevier Ltd (2016). 10.1016/j.neubiorev.2016.06.022

[37] Miller, K.J., Ludvig, E.A., Pezzulo, G., Shenhav, A.: Realigning models of habitual and goal-directed decision-making. In: Goal-Directed Decision Making: Computations and Neural Circuits, (2018). 10.1016/B978-0-12-812098-9.00018-8

[38] Doll, B.B., Simon, D.A., Daw, N.D.: The ubiquity of model-based reinforcement learning (2012). 10.1016/j.conb.2012.08.003

[39] Silva, C., Hare, T.A.: Humans primarily use model-based inference in the two-stage task. Nature Human Behaviour 4(10) (2020) 10.1038/s41562-020-0905-y

[40] Damoiseaux, J.S., Rombouts, S.A.R.B., Barkhof, F., Scheltens, P., Stam, C.J., Smith, S.M., Beckmann, C.F.: Consistent resting-state networks across healthy subjects. Proceedings of the National Academy of Sciences of the United States of America 103(37) (2006) 10.1073/pnas.0601417103

[41] Dastjerdi, M., Foster, B.L., Nasrullah, S., Rauschecker, A.M., Dougherty, R.F., Townsend, J.D., Chang, C., Greicius, M.D., Menon, V., Kennedy, D.P., Parvizi, J.: Differential electrophysiological response during rest, self-referential, and nonself-referential tasks in human posteromedial cortex. Proceedings of the National Academy of Sciences of the United States of America 108(7) (2011) 10.1073/pnas.1017098108

[42] Power, J.D., Cohen, A.L., Nelson, S.M., Wig, G.S., Barnes, K.A., Church, J.A., Vogel, A.C., Laumann, T.O., Miezin, F.M., Schlaggar, B.L., Petersen, S.E.: Functional Network Organization of the Human Brain. Neuron 72(4) (2011) 10.1016/j.neuron.2011.09.006

[43] Raichle, M.E., Raichle, M.E.: Searching for a baseline: Functional imaging and the resting human brain. Nature Reviews Neuroscience 2(10) (2001) 10.1038/35094500

[44] Buckner, R.L., Carroll, D.C.: Self-projection and the brain. Trends in Cognitive Sciences 11(2) (2007) 10.1016/j.tics.2006.11.004

[45] Pearson, J.M., Heilbronner, S.R., Barack, D.L., Hayden, B.Y., Platt, M.L.: Posterior cingulate cortex: Adapting behavior to a changing world (2011). 10.1016/j.tics.2011.02.002

[46] Weber, S., Aleman, A., Hugdahl, K.: Involvement of the default mode network under varying levels of cognitive effort. Scientific Reports 12(1) (2022) 10.1038/s41598-022-10289-7

[47] Roy, M., Shohamy, D., Wager, T.D.: Ventromedial prefrontal-subcortical systems and the generation of affective meaning (2012). 10.1016/j.tics.2012.01.005

[48] Lehmann, K., Bolis, D., Friston, K.J., Schilbach, L., Ramstead, M.J.D., Kanske, P.: An Active-Inference Approach to Second-Person Neuroscience. Perspectives on Psychological Science (2023) 10.1177/17456916231188000

[49] Carhart-Harris, R.L., Friston, K.J.: The default-mode, ego-functions and free-energy: A neurobiological account of Freudian ideas (2010). 10.1093/brain/awq010

[50] Miller, K.J., Botvinick, M.M., Brody, C.D.: Dorsal hippocampus contributes to model-based planning. Nature Neuroscience 20(9), 1269–1276 (2017) 10.1038/nn.4613

[51] Turner, V.S., O’Sullivan, R.O., Kheirbek, M.A.: Linking external stimuli with internal drives: A role for the ventral hippocampus. Elsevier Ltd (2022). 10.1016/j.conb.2022.102590

[52] Corbetta, M., Shulman, G.L.: Control of goal-directed and stimulus-driven attention in the brain. Nature Reviews Neuroscience 3(3) (2002) 10.1038/nrn755

[53] Asplund, C.L., Todd, J.J., Snyder, A.P., Marois, R.: A central role for the lateral prefrontal cortex in goal-directed and stimulus-driven attention. Nature Neuroscience 13(4), 507–512 (2010) 10.1038/nn.2509

[54] Goulden, N., Khusnulina, A., Davis, N.J., Bracewell, R.M., Bokde, A.L., McNulty, J.P., Mullins, P.G.: The salience network is responsible for switching between the default mode network and the central executive network: Replication from DCM. NeuroImage 99 (2014) 10.1016/j.neuroimage.2014.05.052

[55] Corbetta, M., Patel, G., Shulman, G.L.: The Reorienting System of the Human Brain: From Environment to Theory of Mind. Neuron 58(3), 306–324 (2008) 10.1016/j.neuron.2008.04.017

[56] Vossel, S., Geng, J.J., Fink, G.R.: Dorsal and ventral attention systems: Distinct neural circuits but collaborative roles. Neuroscientist 20(2) (2014) 10.1177/1073858413494269

[57] Moutoussis, M., Garzón, B., Neufeld, S., Bach, D.R., Rigoli, F., Goodyer, I., Bullmore, E., Fonagy, P., Jones, P., Hauser, T., Romero-Garcia, R., St Clair, M., Vértes, P., Whitaker, K., Inkster, B., Prabhu, G., Ooi, C., Toseeb, U., Widmer, B., Bhatti, J., Villis, L., Alrumaithi, A., Birt, S., Bowler, A., Cleridou, K., Dadabhoy, H., Davies, E., Firkins, A., Granville, S., Harding, E., Hopkins, A., Isaacs, D., King, J., Kokorikou, D., Maurice, C., McIntosh, C., Memarzia, J., Mills, H., O’Donnell, C., Pantaleone, S., Scott, J., Fearon, P., Suckling, J., Harmelen, A.L., Kievit, R., Guitart-Masip, M., Dolan, R.J.: Decision-making ability, psychopathology, and brain connectivity. Neuron 109(12) (2021) 10.1016/j.neuron.2021.04.019

[58] Huys, Q.J.M., Browning, M., Paulus, M.P., Frank, M.J.: Advances in the computational understanding of mental illness. Springer (2021). 10.1038/s41386-020-0746-4

[59] Rupprechter, S., Stankevicius, A., Huys, Q.J.M., Steele, J.D., Seriés, P.: Major Depression Impairs the Use of Reward Values for Decision-Making. Scientific Reports 8(1) (2018) 10.1038/s41598-018-31730-w

[60] Kim, K., Kim, S.W., Myung, W., Han, C.E., Fava, M., Mischoulon, D., Papakostas, G.I., Seo, S.W., Cho, H., Seong, J.K., Jeon, H.J.: Reduced orbitofrontal-thalamic functional connectivity related to suicidal ideation in patients with major depressive disorder. Scientific Reports 7(1) (2017) 10.1038/s41598-017-15926-0

[61] Cox, R.W.: AFNI: Software for analysis and visualization of functional magnetic resonance neuroimages. Computers and Biomedical Research 29(3) (1996) 10.1006/cbmr.1996.0014

[62] Cox, R.W., Hyde, J.S.: Software tools for analysis and visualization of fMRI data. NMR in Biomedicine 10(4-5) (1997) 10.1002/(SICI)1099-1492(199706/08)10:4/5<171::AID-NBM453>3.0.CO;2-L

[63] Glasser, M.F., Sotiropoulos, S.N., Wilson, J.A., Coalson, T.S., Fischl, B., Andersson, J.L., Xu, J., Jbabdi, S., Webster, M., Polimeni, J.R., Van Essen, D.C., Jenkinson, M.: The minimal preprocessing pipelines for the Human Connectome Project. NeuroImage 80, 105–124 (2013) 10.1016/j.neuroimage.2013.04.127

[64] Dale, A.M., Fischl, B., Sereno, M.I.: Cortical surface-based analysis: I. Segmentation and surface reconstruction. NeuroImage 9(2) (1999) 10.1006/nimg.1998.0395

[65] Fischl, B., Salat, D.H., Busa, E., Albert, M., Dieterich, M., Haselgrove, C., Van Der Kouwe, A., Killiany, R., Kennedy, D., Klaveness, S., Montillo, A., Makris, N., Rosen, B., Dale, A.M.: Whole brain segmentation: Automated labeling of neuroanatomical structures in the human brain. Neuron 33(3) (2002) 10.1016/S0896-6273(02)00569-X

[66] Dickie, E.W., Anticevic, A., Smith, D.E., Coalson, T.S., Manogaran, M., Calarco, N., Viviano, J.D., Glasser, M.F., Van Essen, D.C., Voineskos, A.N.: Ciftify: A framework for surface-based analysis of legacy MR acquisitions. NeuroImage 197 (2019) 10.1016/j.neuroimage.2019.04.078

[67] Robinson, E.C., Jbabdi, S., Glasser, M.F., Andersson, J., Burgess, G.C., Harms, M.P., Smith, S.M., Van Essen, D.C., Jenkinson, M.: MSM: A new flexible framework for multimodal surface matching. NeuroImage 100 (2014) 10.1016/j.neuroimage.2014.05.069

[68] Taylor, P.A., Chen, G., Glen, D.R., Rajendra, J.K., Reynolds, R.C., Cox, R.W.: FMRI processing with AFNI: Some comments and corrections on “Exploring the Impact of Analysis Software on Task fMRI Results”. bioRxiv (2018)

[69] Jo, H.J., Saad, Z.S., Simmons, W.K., Milbury, L.A., Cox, R.W.: Mapping sources of correlation in resting state FMRI, with artifact detection and removal. NeuroImage 52(2) (2010) 10.1016/j.neuroimage.2010.04.246

[70] Schaefer, A., Kong, R., Gordon, E.M., Laumann, T.O., Zuo, X.-N., Holmes, A.J., Eickhoff, S.B., Yeo, B.T.T.: Local-Global Parcellation of the Human Cerebral Cortex from Intrinsic Functional Connectivity MRI. Cerebral Cortex 28(9), 3095– 3114 (2018) 10.1093/cercor/bhx179

[71] Tian, Y., Margulies, D.S., Breakspear, M., Zalesky, A.: Topographic organization of the human subcortex unveiled with functional connectivity gradients. Nature Neuroscience 23(11) (2020) 10.1038/s41593-020-00711-6

[72] Pauli, W.M., Nili, A.N., Michael Tyszka, J.: Data Descriptor: A high-resolution probabilistic in vivo atlas of human subcortical brain nuclei. Scientific Data 5 (2018) 10.1038/sdata.2018.63

[73] Edlow, B.L., Takahashi, E., Wu, O., Benner, T., Dai, G., Bu, L., Grant, P.E., Greer, D.M., Greenberg, S.M., Kinney, H.C., Folkerth, R.D.: Neuroanatomic connectivity of the human ascending arousal system critical to consciousness and its disorders. Journal of Neuropathology and Experimental Neurology 71(6) (2012) 10.1097/NEN.0b013e3182588293

[74] Maretic, H.P., Frossard, P.: Graph Laplacian Mixture Model. IEEE Transactions on Signal and Information Processing over Networks 6, 261–270 (2020) 10.1109/TSIPN.2020.2983139

[75] Friston, K., Schwartenbeck, P., FitzGerald, T., Moutoussis, M., Behrens, T., Dolan, R.J.: The anatomy of choice: Dopamine and decision-making. Philosophical Transactions of the Royal Society B: Biological Sciences 369(1655) (2014) 10.1098/rstb.2013.0481

[76] Smith, R., Friston, K.J., Whyte, C.J.: A step-by-step tutorial on active inference and its application to empirical data. Journal of Mathematical Psychology 107 (2022) 10.1016/j.jmp.2021.102632

[77] Majumdar, G., Yazin, F., Banerjee, A., Roy, D.: Emotion dynamics as hierarchical Bayesian inference in time. Cerebral Cortex 33(7) (2023) 10.1093/cercor/bhac305

