## Supplementary Information for "Individual Differences in Policy Precision: Links to Suicidal Ideation and Network Dynamics" (from another version of this preprint: version 2, 2024-12-11)

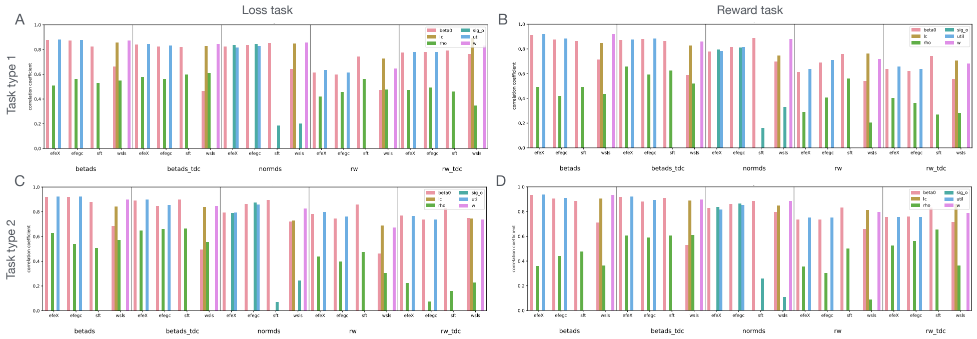


**Fig. 1** Parameter recovery results in Experiment 2

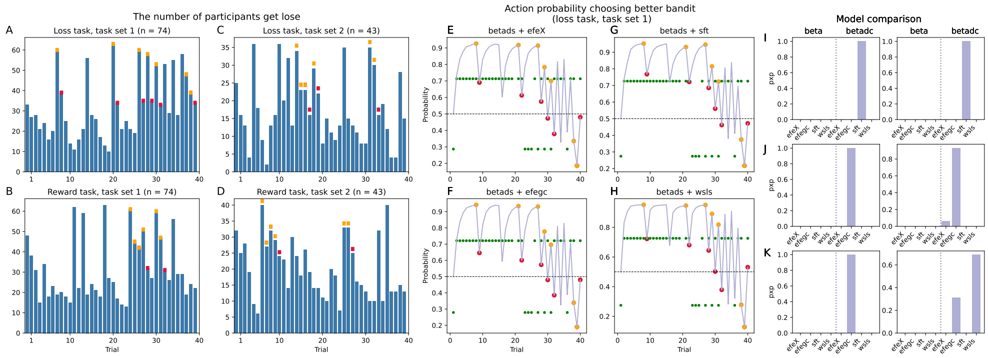


**Fig. 2** Estimation results in Experiment 2

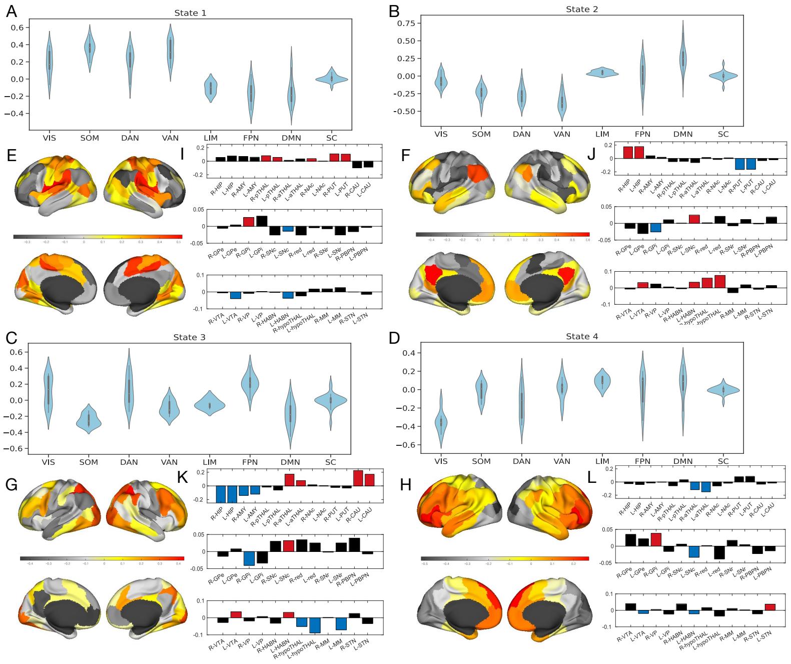


**Fig. 3** Results of the graph Laplacian mixture model (GLMM) analysis. (A) to (H), Mean activation pattern across participants for each brain state estimated from the GLMM. (A) to (D), shows the activation values of regions included in each priori resting state network (RSN). (I) to (L), Mean activations in the subcortical region (SC) of each state. Full names of subcortical region are described in Table1. The regions with significant differences compared with the other states in an analysis of variance (ANOVA) ($p < 0.01$) are colored red if their activation is higher than that of others and blue if lower.

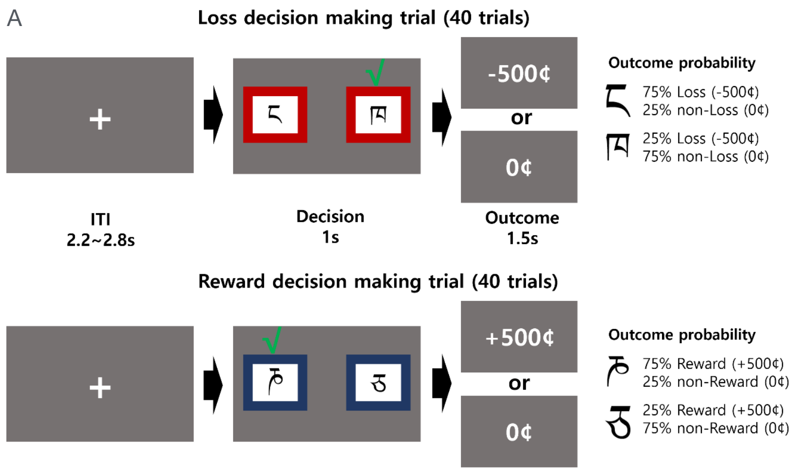


**Fig. 4** Decision-making task used in Experiment 2. (A) In the loss decision-making task trials (choice options with red boundary), choosing one Tibetan character (choice must be made within 0.5 s; green check symbol) caused either loss (-500 ¢ in the Figure) or non-loss (0 ¢ in the Figure), which was presented for 1.5 s. One option resulted in loss with a high probability (75% of loss), while it was highly likely to avoid loss by selecting another option (75% of non-loss). In reward decision-making task trials (choice options with blue boundary), choosing one Tibetan character (green check symbol) caused either reward (+500 ¢ in the Figure) or non-reward (0 ¢ in the Figure). Each option was associated with a high probability of reward (75% of reward) or a high probability of reward omission (75% of non-reward). Inter- trial interval (ITI) varied between 2.2 s and 2.8 s randomly. Note that although we used ¢ in this Figure for understanding, we presented Korean Won for the outcome in the real task.

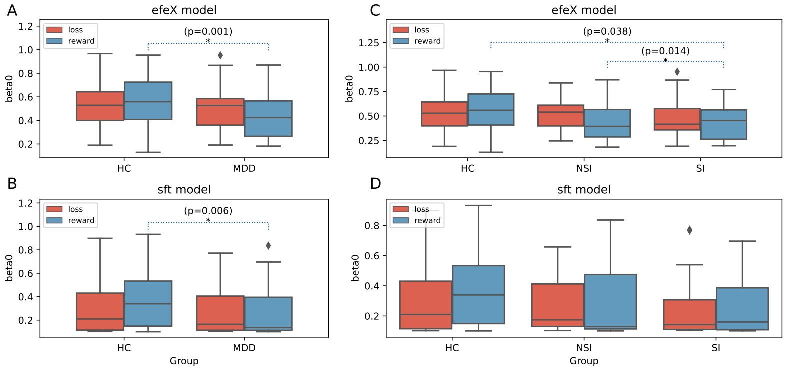


**Fig. 5** ANCOVA results of other models. (A), (B) ANCOVA results comparing *beta0* estimated from *efeX* and softmax models between control (HC) and major depressive disorder (MDD) group. (C), (D) ANCOVA results comparing *beta0* estimated from *efeX* and softmax models between HC and MDD without suicidal ideation (NSI) and with suicidal ideation (SI) group. Asterisks (*) denotes statistical significance in ANCOVA.

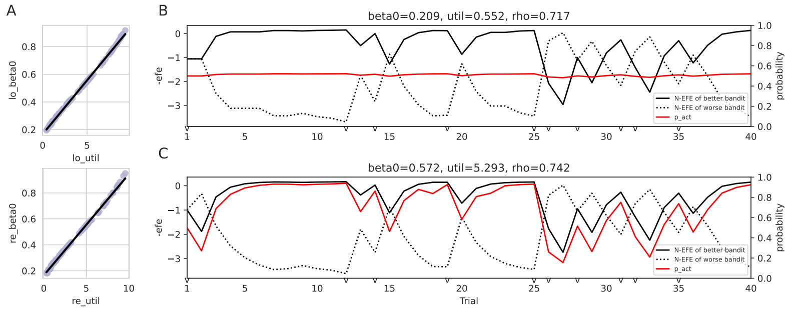


**Fig. 6** (A) High correlations between the *beta0* and *util* parameters in both tasks. (B), (C) The example trajectories of minus expected free energy (N-EFE) of both bandits and action probability of choosing better bandit (p_act) along the trials in low and high *beta0* participants, respectively. In active inference framework, participants choose the bandit with lower EFE (lager N-EFE). Therefore, as the difference in EFE between the two choices increases, the probability of choosing the better bandit also increases. And the *beta0* values for each individual determine the extent to which the EFE (black line) influences the calculation of action probability (red line). As *beta0* decreases, the impact of EFE on action probability diminishes. In such cases, the action probabilities of tend to hover around 0.5 with low *beta0*.

**Table 1** Result of ANOVA comparing the activations of each subcortical region across states

| Subcortex | Sum of Squares | F | Sig. | Post-hoc |
|---|---|---|---|---|
| 1. Rt. Hippocampus | 5.4895 | 74.65 | < 0.0001 | State2 > State 1, 4 > State 3 |
| 2, Lt. Hippocampus | 6.7703 | 99.74 | < 0.0001 | State2> State 1> State 4> State 3 |
| 3. Rt. Amygdala | 1.4059 | 20.56 | < 0.0001 | State1, 2, 4> State 3 |
| 4. Lt. Amygdala | .9968 | 19.14 | < 0.0001 | State1, 2, 4> State 3 |
| 5. Rt. posterior thalamus | .6658 | 11.62 | < 0.0001 | State1> State 2, 3, 4 |
| 6. Lt. posterior thalamus | .5348 | 9.53 | < 0.0001 | State 1, 4> State 2, 3 |
| 7. Rt. anterior thalamus | 2.4144 | 35.79 | < 0.0001 | State 3> State 1> State 2, 4 |
| 8. Lt. anterior thalamus | 1.5685 | 20.75 | < 0.0001 | State 1, 2, 3> State 4 |
| 9. Rt. nucleus accumbens | .3187 | 5.58 | .0011 | State 1, 3> State 4 |
| 10. Lt. nucleus accumbens | .0121 | 0.27 | .8491 | - |
| 11. Rt. putamen | 2.6004 | 34.21 | < 0.0001 | State1, 4> State 3> State 2 |
| 12. Lt. putamen | 2.6540 | 39.53 | < 0.0001 | State1, 4> State 3> State 2 |
| 13. Rt. caudate | 3.3277 | 49.44 | < 0.0001 | State 3> State 1, 2, 4 |
| 14. Lt. caudate | 1.9577 | 28.79 | < 0.0001 | State 3> State 1, 2, 4 |
| 15. Rt. GPe | .1174 | 3.25 | .0227 | - |
| 16. Lt. GPe | .0677 | 1.56 | .1993 | - |
| 17. Rt. GPi | .1925 | 3.98 | .0088 | State 1, 4> State 3 |
| 18. Lt. GPi | .1355 | 2.87 | .0372 | - |
| 19. Rt. SNc | .0877 | 2.09 | .1028 | - |
| 20. Lt. SNc | .1659 | 3.94 | .0092 | State 2,3> State 4 |
| 21. Rt. red nucleus | 0.11 | 2.57 | .0549 | - |
| 22. Lt. red nucleus | .1347 | 3.48 | .0168 | - |
| 23. Rt. SNr | .0223 | 0.52 | .6656 | - |
| 24. Lt. SNr | .0808 | 1.65 | .1779 | - |
| 25. Rt. parabrachial pigmented nucleus | .1349 | 3.18 | .0249 | - |
| 26. Lt. parabrachial pigmented nucleus | .0456 | 1.15 | .3295 | - |
| 27. Rt. VTA | .1604 | 3.56 | .0151 | - |
| 28. Lt. VTA | .2252 | 4.74 | .0032 | State 2,3> State 1,4 |
| 29. Rt. ventral pallidum | .0473 | 0.96 | .4107 | - |
| 30. Lt. ventral pallidum | .0375 | 0.68 | .5625 | - |
| 31. Rt. habenula | .1648 | 3.49 | .0167 | - |
| 32. Lt. habenula | .2137 | 4.45 | .0047 | State 2,3> State 1,4 |
| 33. Rt. hypothalamus | .3529 | 8.11 | < 0.0001 | State 2> State 1, 3 |
| 34. Lt. hypothalamus | .7498 | 16.73 | < 0.0001 | State 2> State 3, 4<br>State 1> State 3 |
| 35. Rt. mammillary body | .0515 | 1.04 | .375 | - |
| 36. Lt. mammillary body | .3342 | 7.08 | .0001 | State 1, 2, 4> State 3 |
| 37. Rt. subthalamic nucleus | .0802 | 1.64 | .1813 | - |
| 38. Lt. subthalamic nucleus | .1879 | 4.5 | .0044 | State 4> State 3 |
| 39. dorsal raphe | .1328 | 2.99 | .032 | - |
| 40. locus coeruleus | .1009 | 2.07 | .1059 | - |
| 41. lateraldorsal tegmental nucleus | .0978 | 2.19 | .0897 | - |
| 42. median raphe | .0510 | 1.26 | .2877 | - |
| 43. mesencephalic reticular formation | .1234 | 2.92 | .035 | - |
| 44. periaqueductal grey | .0245 | 0.47 | .7036 | - |
| 45. parabrachial complex | .1445 | 3.27 | .022 | - |
| 46. pontis oralis | .0482 | 1.12 | .3432 | - |
| 47. pedunculotegmental nucleus | .1311 | 2.99 | .0321 | - |
